# Single-cell Transcriptome and Accessible Chromatin Dynamics During Endocrine Pancreas Development

**DOI:** 10.1101/2022.01.28.478217

**Authors:** Eliza Duvall, Cecil M. Benitez, Krissie Tellez, Martin Enge, Philip T. Pauerstein, Lingyu Li, Songjoon Baek, Stephen R. Quake, Jason P. Smith, Nathan C. Sheffield, Seung K. Kim, H. Efsun Arda

## Abstract

Delineating gene regulatory networks that orchestrate cell-type specification is an ongoing challenge for developmental biology studies. Single-cell analyses offer opportunities to address these challenges and accelerate discovery of rare cell lineage relationships and mechanisms underlying hierarchical lineage decisions. Here, we describe the molecular analysis of pancreatic endocrine cell differentiation using single-cell gene expression, chromatin accessibility assays coupled to genetic labeling and cell sorting. We uncover transcription factor networks that delineate *β*-, *α*- and *δ*-cell lineages. Through genomic footprint analysis we identify transcription factor-regulatory DNA interactions governing pancreatic cell development at unprecedented resolution. Our analysis suggests that the transcription factor Neurog3 may act as a pioneer transcription factor to specify the pancreatic endocrine lineage. These findings could improve protocols to generate replacement endocrine cells from renewable sources, like stem cells, for diabetes therapy.

## INTRODUCTION

More than 400 million people are living with diabetes worldwide. Diabetes results from loss or dysfunction of hormone-producing endocrine islet cells in the pancreas, whose principal role is to regulate circulating glucose levels. Recent advances in tissue engineering to replace non-functioning endocrine cells have renewed interest in understanding the molecular mechanisms of pancreatic endocrine cell differentiation (Siehler et al., 2021).

A key event during endocrine pancreas development is expression of the transcription factor Neurog3 in select pancreatic duct cells (Gradwohl et al., 2000). Neurog3 specifies endocrine progenitor cells, which differentiate into hormone producing cells that delaminate from the duct and aggregate to form pancreatic islets (reviewed in Arda et al., 2013; Bastidas-Ponce et al., 2017; Benitez et al., 2012). Several distinct endocrine cell types aggregate within pancreatic islets, including insulin^pos^ *β*-cells, glucagon^pos^ *α*-cells, somatostatin^pos^ *δ*-cells, ghrelin^pos^ *ε*-cells, and PPY^pos^ *γ*-cells. Mice lacking pancreatic *Neurog3* fail to develop endocrine islet cells (Gradwohl et al., 2000; Gu et al., 2002; Schwitzgebel et al., 2000; Smith et al., 2004). In one model based on lineage tracing (Kopinke et al., 2011; Solar et al., 2009), Neurog3^pos^ cells originate from a “bi-potent progenitor” with potential to generate either ducts or islets (reviewed in Bankaitis et al., 2015).

Emerging single-cell technologies are revolutionizing developmental biology by enabling quantitative molecular analysis of transient, rare cell types in developing organs, especially lineage progenitor cells. Recently, several groups used single-cell RNA sequencing (scRNA-Seq) to catalog dynamic transcriptome changes during mouse pancreas development and endocrine cell differentiation (Bastidas-Ponce et al., 2019; Byrnes et al., 2018; Krentz et al., 2018; Qiu et al., 2017a; Sharon et al., 2019; Yu et al., 2019). Some studies suggested that endocrine progenitor subtypes exist or are biased towards specific hormone lineages (Liu et al., 2019; Scavuzzo et al., 2018; Yu et al., 2019). While these reports contributed substantially to our understanding of endocrine pancreas development, no study has yet reported specification of the crucial islet *δ*-cell lineage (Arrojo e Drigo et al., 2019), or investigated chromatin conformation changes by overcoming cell labeling ambiguities related to *Neurog3*-GFP cells (Lee et al., 2002).

To address these unmet needs, we used an integrative approach that combined cell surface marker-based sorting, genetic labeling, chromatin analysis, and single-cell assays to elucidate molecular mechanisms underlying gene expression changes during endocrine pancreas differentiation. By establishing pseudotime trajectories for hormone lineages, including islet *δ*-cells, we identified unique combinations of transcription factors guiding differentiation of the *β*-, *α*-, and *δ*-lineages. Chromatin accessibility analysis using ATAC-seq unexpectedly revealed extensive similarities between duct cells and those that activate *Neurog3*. We discovered genomic regions that undergo substantial transformation during development and identified enriched motifs in open chromatin specific to differentiation stages. We also applied powerful genomic footprint analysis to identify transcription factor activity in open chromatin regions and found evidence of specific transcription factor footprints linked to their associated motifs. Our analysis suggests a revised model for endocrine pancreas development by providing evidence for direct development of this lineage from duct cells, and the absence of a bipotent progenitor.

Our results demonstrate the feasibility of using a combined scRNA-seq and ATAC-seq analysis to map gene regulatory networks that define pancreatic cell lineages. We anticipate our findings and those from similar work should foster efforts aiming to direct development of renewable cell sources, like stem cells, for tissue replacement and regeneration.

## RESULTS

### Single-cell transcriptomic analysis of endocrine pancreas development

To understand gene expression dynamics during pancreatic endocrine cell differentiation, we performed scRNA-Seq on cells isolated from mouse embryonic day 15.5 (E15.5) and E17.5 pancreas. We used the *Neurog3*-*eGFP* knock-in and *Neurog3-Cre,Rosa-mTmG* mice combined with cell surface markers to isolate specific populations from the embryonic pancreas (see **Methods**) (Lee et al., 2002; Muzumdar et al., 2007; Sugiyama et al., 2007). We followed the Smart-Seq2 protocol to sequence mRNAs from single-cells sorted into 96-well plates by fluorescence-activated cell sorting (FACS, **Supplementary Figure 1A**). Using this strategy, we collected and sequenced a total of 604 cells: 461 from E15.5 cells and 143 from E17.5 cells.

After initial read processing to count transcripts for each gene in each cell (**Supplementary Table 1**), we used Monocle2— a single-cell analysis tool, for downstream cell clustering and trajectory analysis (**Supplementary Figure 2**). Unsupervised clustering organized cells based on transcriptome similarity, revealing a recognizable sequence of pancreatic endocrine cell differentiation (**Figure 1A, Supplementary Figure 1B**). This developmental process included a progenitor cluster expressing high levels of *Neurog3*, a transitioning, early endocrine cell cluster, a definitive endocrine cluster marked by high levels of *Chga* expression, and a cluster of exocrine cells marked by *Cpa1* expression (**Figure 1B**). We also found a small cluster of mesenchymal cells (14 cells, < 3% of total cells), which were excluded from further analysis.

**Figure 1:**
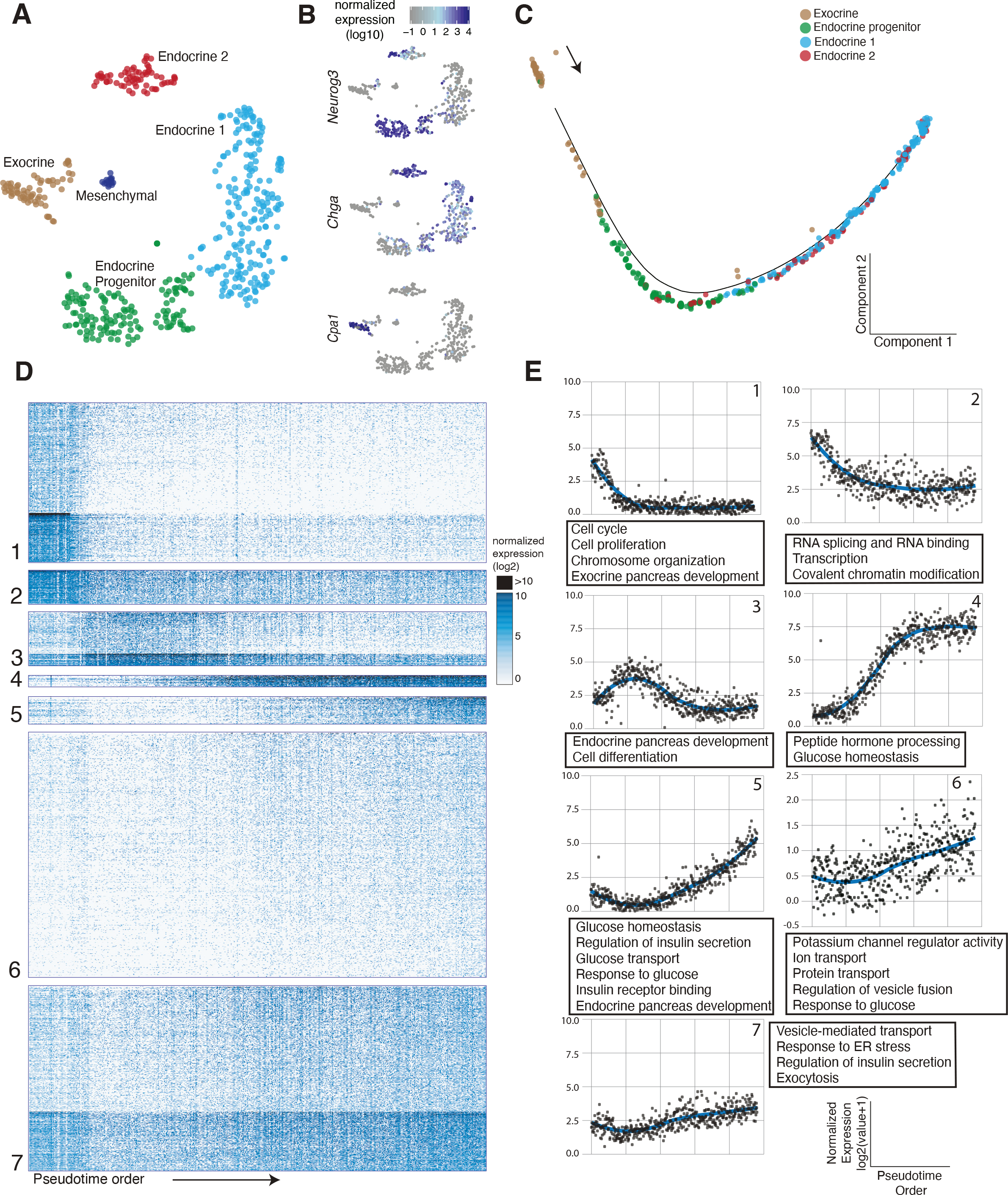
(A) t-SNE plot showing single-cell clusters, colored by cluster. Each dot is a single-cell. Cluster names are indicated on the graph. (B) Marker gene expression levels overlaid onto the *t-*SNE plot. *Cpa1* (exocrine), *Neurog3* (endocrine progenitor), and *Chga* (pan-endocrine). (C) Alignment of single-cells onto a pseudotime trajectory beginning from duct cells and ending with hormone producing endocrine cells. Colors represent the clusters in (A). (D) Heat map representation of > 2500 differentially expressed genes during pancreatic endocrine cell differentiation, organized into different clusters. Rows represent genes, columns represent single-cells ordered by the pseudotime order. (E) Graphs representing expression trends per cluster determined by fitting a loess curve of average gene expression per cluster, plotted over pseudotime. Each point represents the average expression of genes within each cluster for a single-cell along pseudotime. Associated GO terms are listed in the text boxes.

To delineate gene expression programs involved in endocrine cell development, we aligned cells in a pseudotime trajectory based on quantitative gene expression profiles that change continuously in differentiating cells. This analysis placed all cells in a single trajectory that corroborated the known progression of duct cells into *Neurog3^pos^* progenitors, followed by hormone expressing endocrine cells (**Figure 1C**). We found more than 2,500 genes whose expression changed significantly along this pseudotime trajectory (*q*-value < 0.05). *k-*means analysis partitioned these differentially expressed genes into distinct gene clusters (**Figure 1D, Supplementary Table 2**). To better visualize the gene expression trends in each cluster, we used LOESS smoothing along pseudotime (**Figure 1E**; **Methods**). GO term analysis identified enriched biological process terms in these clusters relevant to pancreatic differentiation (FDR < 0.2, **Figure 1E**, **Supplementary Table 3**; (Arda et al., 2013; Bastidas-Ponce et al., 2017).

**Cluster 1** included genes that are expressed at high levels at the start of the pseudotime trajectory, then decline significantly or are extinguished as cells differentiate into the endocrine lineages. These genes included known regulators of multipotent pancreatic progenitor or exocrine cells (*Ptf1a*, *Hes1*, *Notch1*, *Rbpj*), the cell cycle (*Mki67*, *Ccna2*, *Cdk1*), and factors involved in maintenance of chromosome organization or covalent chromatin modifications (*Smc4*, *Ezh2* and *Ctcf)*. **Cluster 2** genes had a similar trend, although their expression remained detectable in endocrine cells. These include genes regulating RNA binding and splicing, translation initiation, and ribonucleoprotein complexes. **Cluster 3** genes are mainly expressed in endocrine progenitor cells and trending similarly with *Neurog3* expression, including *Pax4*, *Tox3*, and *Cbfa2t3*. Most Cluster 3 transcripts were only detectable transiently in progenitor cells, then extinguished in endocrine cells. Cluster 3 was associated with GO terms related to cell differentiation and endocrine pancreas development (**Supplementary Table 3**).

Clusters 4-6 contained genes whose expression increased following the *Neurog3* induction. **Cluster 4** genes included *Chga*, *Pcsk2*, *Pax6*, *Iapp*, *Neurod1* and *Isl1*, and were turned on shortly after *Neurog3* expression peaked, in early endocrine cells that still lack mRNAs encoding the principal islet hormones. **Clusters 5 and 6** genes include the hormones, *Ins1*, *Ins2, Ppy, Sst* and *Gcg*, whose expression peak in endocrine cells. These clusters also included genes involved in vesicle mediated transport, ion transport, response to ER stress, regulation of insulin secretion, and exocytosis. **Cluster 7** contains genes enriched with functions in the mitochondrial respiratory chain complex, proton transport, and ATP synthesis. Taken together, pancreatic endocrine cell specification involves highly dynamic gene regulatory programs, multiple groups of gene families with distinct functions.

### Analysis of pancreatic endocrine progenitors

Prior studies reported the existence of distinct *Neurog3^pos^* endocrine progenitor subtypes (Liu et al., 2019; Scavuzzo et al., 2018; Yu et al., 2019). To investigate the heterogeneity in *Neurog3^pos^* progenitor cells, we focused on the cells expressing *Neurog3* transcript in our dataset and visualized them using the *t-*SNE method. This analysis identified three clusters based on *Neurog3* transcript abundance— designated as *high*, *medium* and *low*, though none of the clusters split into visually distinct groups on the *t*-SNE projection (**Figure 2A**). The *Neurog3^hi^* cells had the highest *Neurog3* levels compared to other clusters (**Figure 2B**), likely the result of increased *Neurog3* transcription that occurs during the secondary transition of endocrine differentiation (Schwitzgebel et al., 2000). Less than 10% of the *Neurog3^hi^* cells had detectable *Chga* expression (**Figure 2C**). In *Neurog3^med^* and *Neurog3^lo^* cells *Neurog3* transcript levels decreased, while *Chga* levels increased (**Figure 2C**). Thus, the observed ‘transcriptional heterogeneity’ in *Neurog3^pos^* cells is a direct reflection of advancing development. Moreover, this data argues against a model where endocrine progenitor cells randomly develop from cells with heterogeneous *Neurog3* levels. When we analyzed the expression of individual hormone genes, we found that the number of cells expressing *Ins1, Ins2*, *Gcg*, or *Sst* increased as cells transitioned from *Neurog3^hi^* to *Neurog3^lo^* progenitors, with *Sst* appearing only in the *Neurog3^lo^* cluster (**Figure 2D**). Additionally, we investigated the number of cells simultaneously expressing one, two, or three of these hormone genes and found that the number of cells co-expressing multiple hormone genes increased as *Neurog3* expression decreases. For instance, none of the *Neurog3^hi^* cells were polyhormonal, whereas 18% of *Neurog3^lo^* cells expressed two, and 2% expressed all three hormone genes (**Figure 2E**).

**Figure 2:**
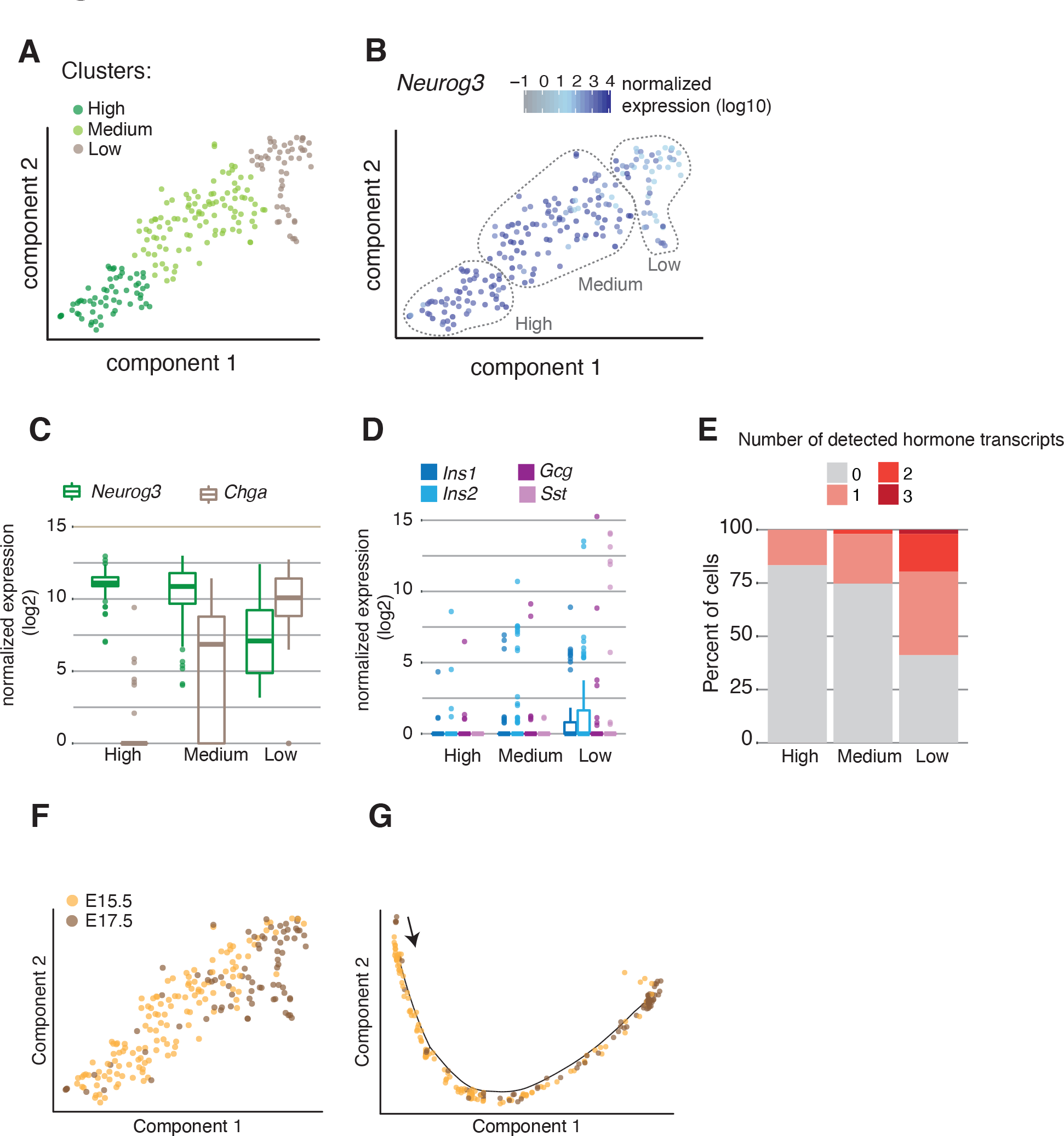
(A) t-SNE plot showing *Neurog3* expressing cell subsets. Each dot is a single-cell, colored by clusters, or (B) *Neurog3* expression. (C) Box plots show normalized *Neurog3* and *Chga* expression in each cluster. (D) Box plots show normalized hormone transcripts detected in each cluster. (E) Stacked bar plot showing the percent of cells within each cluster expressing zero, one, two, or three hormone genes (*Ins1* or *Ins2, Gcg, Sst*). (F) t-SNE projection of *Neurog3* expressing cells colored by the embryonic day they were isolated. (G) Pseudotime trajectory of *Neurog3* positive cells colored by the embryonic day they were isolated.

To investigate whether there is transcriptional heterogeneity in *Neurog3^pos^* endocrine progenitors isolated from different developmental stages, we examined all *Neurog3^pos^* cells by incorporating the embryonic stage information onto the clusters (**Figure 2F**). We did not observe distinct clustering of E15.5 and E17.5 *Neurog3^pos^* endocrine progenitors; rather, the cells were arranged coincident with their developmental stage (**Figure 2F**). When temporally ordering *Neurog3^pos^* cells via pseudotime analysis, the continuous developmental progression was apparent in a single trajectory, without any branching (**Figure 2G**). Taken together, in our dataset we did not find evidence for lineage biases or subtypes in endocrine progenitors isolated from different embryonic time points. We found that nascent endocrine cells may transiently co-express mRNAs encoding multiple hormones in an intermediate ‘polyhormonal’ state preceding branch specification.

### Single-cell trajectories defining endocrine cell type specification

While *Neurog3* is necessary and sufficient to establish the pancreatic endocrine lineage, the mechanisms underlying subsequent endocrine lineage diversification are not well established. Other studies using single-cell approaches successfully delineated *β*- and *α*-cell branches of islet endocrine cell differentiation, but failed to identify a clear branch for *δ*-cell specification (Byrnes et al., 2018; Liu et al., 2019; Qiu et al., 2017a; Scavuzzo et al., 2018; Sharon et al., 2019; Yu et al., 2019). In our data, an unsupervised approach including all cells also did not yield to trajectories defining individual hormone lineages (**Figure 1C**). We reasoned that when all cells are included, the substantial change in gene expression programs at the onset of *Neurog3* activation might hinder the discovery of less pronounced differences in the initial *β*-, *α*-, and *δ*-cell lineage decisions. To circumvent this issue, we focused analysis on cells after *Neurog3* peak expression (**Supplementary Figure 3**) and performed semi-supervised clustering with marker gene information (Qiu et al., 2017b). Briefly, endocrine progenitors, *β*-, *α*-, and *δ*-cells were pre-assigned based on marker genes before attempting clustering. A prior study used a similar approach to resolve mixed hematopoietic lineages (Iterative Clustering and Guide-gene Selection, Olsson et al., 2016). We then performed iterative rounds of trajectory analysis, sequentially removing cells already assigned to an endocrine cell branch in each iteration, until all branches were identified (**Figures 3A-B**). This approach successfully partitioned *β*-, *α*- and *δ*-cells into nearly exclusive, specific branches (**Figure 3C**) suggesting that expert curation can overcome some limitations of trajectory analysis (also see **Discussion**).

**Figure 3:**
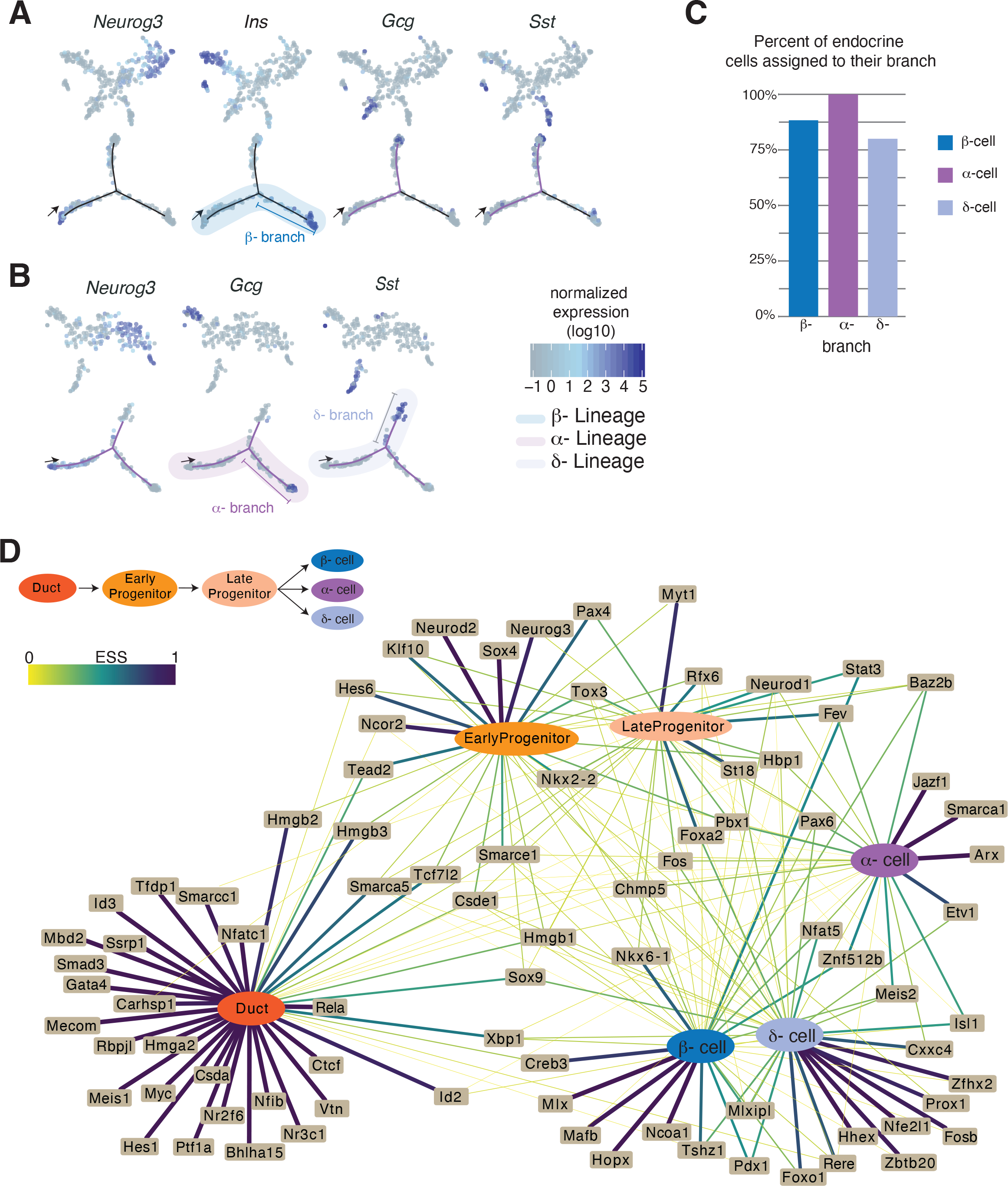
(A) (top) t-SNE plots showing semi-supervised clustering of single cells, first iteration to resolve *β*-lineage. Each dot is a single cell, colored by marker gene expression. (bottom) Trajectory of cells beginning at the arrow, each dot is a single cell and are colored by marker gene expression. High expression of *Ins1* and *Ins2* is seen in cells at the end of the *β*-branch. (B) (top) *t*-SNE plots showing semi-supervised clustering of single cells, second iteration to resolve the *α*- and *δ*-lineages. Each dot is a single cell, colored by marker gene expression. (bottom) Trajectory of cells beginning at the arrow, each dot is a single cell and are colored by marker gene expression. High expression of *Gcg* is seen on the *α*- branch, and *Sst* is seen in cells at the end of the *δ*-branch. (C) Bar graph indicating the percent of endocrine cells that were assigned in the appropriate branch; 88% for *β*-, 100% for *α*-, and 80% for *δ*-cells (D) Network showing the relationship between TF expression and cell state. The edges represent the expression specificity of TFs in each state. Thickness and color of the edges directly correspond to the expression specificity scores (ESS, see **Methods**). ESS values range from 0 to 1, where ESS=1 means TF is exclusively expressed in that cell type, and ESS=0 means no expression. Ubiquitous expression is ESS=0.166.

### TF networks regulating islet cell lineage gene expression

To reveal the gene expression changes underlying distinct trajectories of endocrine cell specification, we performed differential gene expression analysis between cells assigned to the *β*-, *α*- and *δ*-lineages. We defined the lineages as beginning from the duct cells and ending with hormone expressing endocrine cells (**Figures 3A-B**, **Supplementary Table 4**). We focused our analysis on transcription factors (TFs) due to their well-established role in determining cell fates. This analysis revealed 145 TFs whose expression changed significantly during endocrine cell differentiation (**Supplementary Figure 4**). We visualized how these TFs may be regulating distinct lineages by constructing a network based on TF expression patterns in each cell type (duct, *β*-, *α*- and *δ*-cells) or state (early progenitor, late progenitor; **Supplementary Table 5**, also see **Methods** for details). For instance, *Hes1* was detected in duct cells, and thus was connected to the node representing the duct cell.

Topological examination of the TF expression-cell state interaction network revealed three network patterns. In **Network Pattern 1**, we found TFs highly specific to a single lineage. For example, 92% of cells in the *β*-cell lineage express *Nkx6-1* and 71% of *α*-cells express *Arx*.

Nkx6-1 is thought to repress transcription of *Arx*, which specifies the *α*-cell lineage; conversely, Arx is postulated to repress transcription of *Nkx6-1*, which specifies the *β*-cell lineage (Schaffer et al., 2013). We found that *Smarca1* is highly specific to the *α*-cell lineage, and this is consistent with recent reports of *Smarca1* activation during *α*-cell development, prior to *Gcg* expression (Byrnes et al., 2018; Yu et al., 2019). *Smarca1* is an ATP-dependent chromatin remodeler, which can be selectively recruited to cell type-specific enhancer elements (Vierbuchen et al., 2017). A second TF, *Etv1* is a Neurog3 target, (Benitez et al., 2014) and in our data we find *Etv1* is highly specific to the fetal *α*-cell lineage indicating this TF has a functional role in *α*-cell development. In our network, we confirmed that *Hhex* is specific to the *δ*- lineage (Zhang et al., 2014), and found additional factors. *Zbtb20* has increased expression in *δ*- cells relative to *β*- and *α*-cells and to our knowledge, has not been reported before. Instead, *Zbtb20* was recently identified as a TF upregulated in the *α*-cell lineage (Yu et al., 2019).

Because the *δ*- lineage was not defined in this report, it is possible that the uncategorized *δ*-cells aligned with the *α*-lineage instead. Other TFs that are highly specific to the *δ*-cell lineage but with no known functions include *Zfhx2*, *Rere*, and *Cxxc4*.

In **Network Pattern 2**, we found TFs that are expressed in multiple cell types or states. For instance, the high mobility group proteins *Hmgb2*, *Hmgb3*, and *Tead2*, a YAP signaling factor, are initially expressed in duct cells, and continue to be expressed in early *Neurog3^pos^* progenitors. We also found known TFs, including *Isl1*, *Rfx6*, *Pax6,* and *Meis2* in the *β*-, *α*-, and *δ*-cell lineages. In line with a prior report, almost all endocrine cells in the *β*-, *α*-, and *δ*- lineages appear to pass through a *Fev*^pos^ stage after *Neurog3* expression (Byrnes et al., 2018). In this network, *Fev* is most specific to late progenitors. After islet cells transit through a *Fev*^pos^ stage, *Fev* expression rapidly declines in the *β*-cell lineage but remains at detectable levels in *α*- and *δ*- cells (**Supplementary Figure 4**).

**Network Pattern 3** includes TFs that follow an ON-OFF-ON pattern as cells differentiate from duct to progenitors to endocrine lineages. For example, *Xbp1* is abundant in duct cells, but its levels decrease in early and late *Neurog3^pos^* progenitors, then increases in *β*-, *α*-, and *δ*-cells. In mice, loss of *Xbp1* results in hyperglycemia (Lee et al., 2011), abnormal zymogen granules and aplasia of acinar cells (Hess et al., 2011). *Xbp1* is an essential regulator of the unfolded protein response and endoplasmic reticulum (ER) stress (reviewed in Hetz, 2012). Similarly, *Creb3* and *Id2* follow the ON-OFF-ON pattern. These TFs were recently reported to be associated with ER and oxidative stress response programs in human islet *β*-cells (Xin et al., 2018).

### Chromatin accessibility dynamics during islet endocrine cell differentiation

To investigate chromatin accessibility changes during endocrine cell differentiation, we performed ATAC-seq (Buenrostro et al., 2013) on purified populations of duct, endocrine progenitor, and endocrine cells isolated from E15.5 pancreas using the *Neurog3*-eGFP knock-in mice (Lee et al., 2002) (**Figure 4A, Supplementary Table 6**). In these mice, the coding region of *Neurog3* is replaced by an eGFP cassette, thereby regulating eGFP production from the endogenous *Neurog3 cis-*regulatory element, including the promoter. As reported previously, heterozygous *Neurog3^eGFP/+^* animals form a complete endocrine pancreas with no discernable phenotypes (Lee et al., 2002). However, in homozygous *Neurog3^eGFP/eGFP^* animals, eGFP^pos^ cells lack Neurog3 and fail to differentiate further into the endocrine lineage.

**Figure 4:**
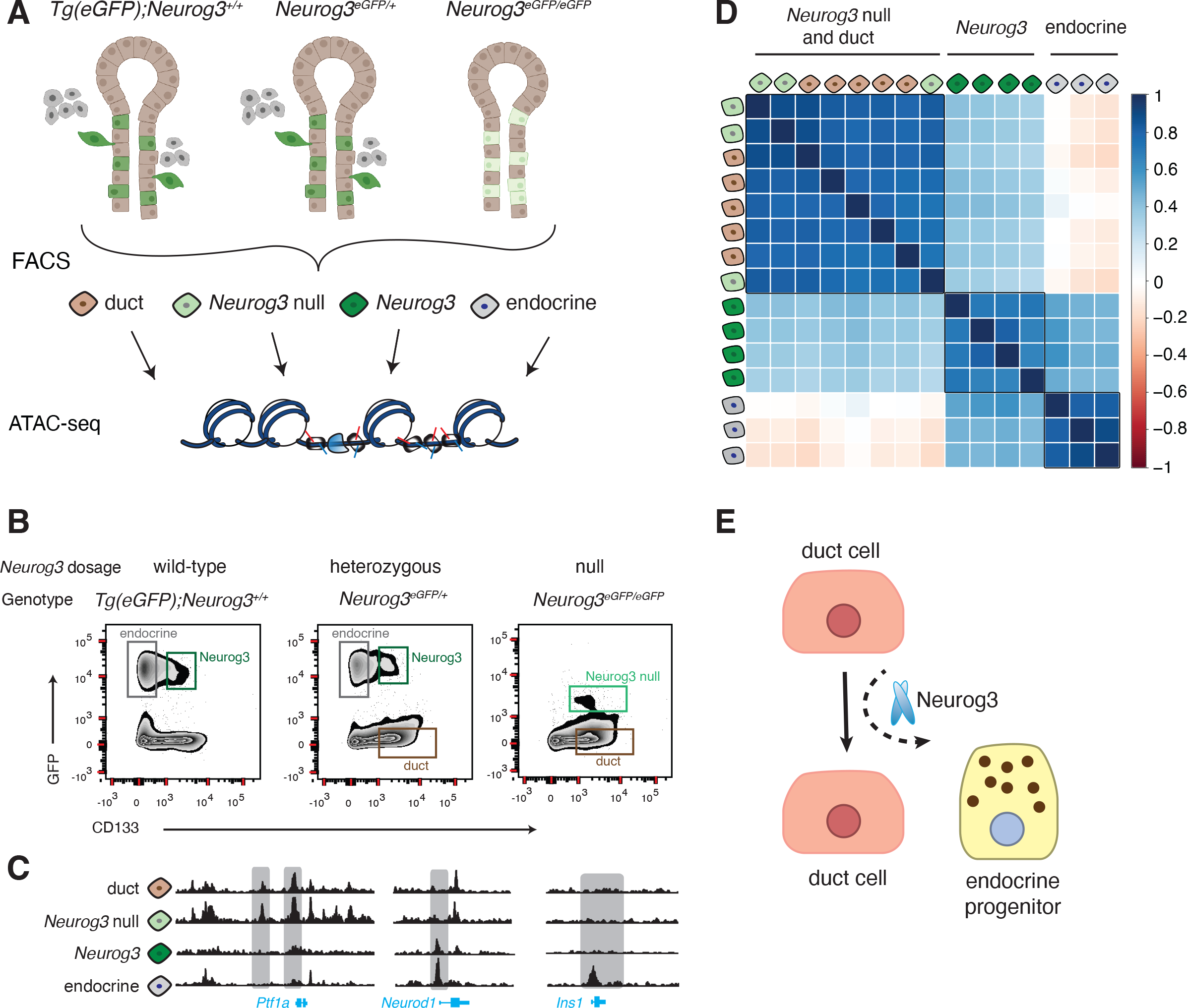
(A) ATAC-seq workflow used in this study. (B) Representative FACS plots showing sorted cell populations and gating strategy. Three mouse genotypes were used to collect four types of cell populations from E15.5 embryos. (C) ATAC-seq reads obtained from different cell populations visualized on the UCSC browser near the three gene loci; *Ptf1a*, *Neurod1*, *Ins1*. (D) Pearson correlation matrix showing the similarity of ATAC-seq samples. Value of 1 indicates high correlation, 0 indicates no correlation, and -1 indicates anti-correlation. The samples are colored as in (A). (E) Suggested model for endocrine pancreas differentiation, where Neurog3 functions as a pioneer factor to shift the default ductal lineage to the endocrine lineage.

To achieve requisite specificity needed for experiments involving purification of *Neurog3-* expressing cells, we managed two concerns not addressed in prior studies (Scavuzzo et al., 2018; Xu et al., 2014). First, since Neurog3 protein stability is transient and short-lived compared to eGFP (White et al., 2008), we needed methods to discriminate between eGFP^pos^ Neurog3^pos^ progenitors and eGFP^pos^ Neurog3^neg^ endocrine cells that have ceased to express *Neurog3*. We achieved this using modified cell sorting strategies (Sugiyama et al., 2007; see **Methods**). Second, to address possible concerns about *Neurog3* gene dosage effects on endocrine cell differentiation, we used mice that are wild-type (“*Tg(eGFP)*; *Neurog3*”, Gu et al., 2004), heterozygous, or homozygous null for *Neurog3* (**Figure 4B**). This enabled direct comparison of chromatin states in endocrine progenitor cells with varying *Neurog3* gene dosage. Specifically, we analyzed four distinct cell populations in different genetic backgrounds: (1) Neurog3^pos^ hormone^neg^ cells (Neurog3); (2) eGFP^pos^ *Neurog3-*null cells (Neurog3 null); (3) hormone^pos^ islet cells (endocrine); and (4) duct cells (duct), (**Figures 4A-B**; **Supplementary Table 6**). In total we performed ATAC-seq on 15 primary pancreatic cell samples.

After aligning sequencing reads, we visually inspected loci near genes essential for pancreas development like *Ptf1a*, *Neurod1* and *Ins1* (**Figure 4C**). ATAC-seq revealed substantial reorganization of chromatin accessibility in regions near these and other genes (see below) during differentiation from duct cells to *Neurog3*^pos^ endocrine progenitor cells, and endocrine cells. For instance, open chromatin “control regions” in the *Ptf1a* locus were detected in wild-type duct cells and *Neurog3*-null cells; the accessibility of this chromatin was then eliminated as duct cells transitioned into endocrine progenitors, a ‘closed’ state also maintained in endocrine cells (Masui et al., 2008). In *Neurod1*, an established Neurog3 target, promoter-proximal chromatin was closed in duct cells but became accessible in Neurog3^pos^ endocrine progenitors. In the *Ins1* locus, chromatin in control regions remained closed until cells committed to the endocrine lineage. Thus, cell purification combined with ATAC-seq generated chromatin maps that corresponded to distinct differentiation stages.

To investigate the similarity in chromatin states between ATAC-seq samples, we calculated pairwise Pearson correlation coefficients and organized samples by clustering (**Figure 4D**). This analysis revealed three groups that corresponded to duct cells, Neurog3^pos^ progenitors and endocrine cells. Chromatin profiles of cells isolated either from wild-type or heterozygous *Neurog3* mice were similar. Unexpectedly, *Neurog3*-null cells clustered with wild-type duct cells (**Figure 4D**). If ductal epithelia harbored bipotent cells that could become either endocrine progenitors or duct cells, we expected to see a *distinct* clustering of Neurog3-null from duct cells. Thus, cells that activated *Neurog3* transcription in the ductal epithelium, but could not differentiate into endocrine lineage have chromatin that is indistinguishable from duct cells. This suggests that chromatin ‘priming’ in duct cells prior to expression of *Neurog3* is not required for endocrine differentiation. Furthermore, Neurog3 might be a *pioneer* transcription factor, whose functions include the capacity to initiate nucleosome displacement or conformational changes in inaccessible chromatin (**Figure 4E**) (Zaret and Mango, 2016).

### Differentially accessible chromatin regions reveal *cis*-regulatory elements that mediate endocrine lineage specification

To identify differentially accessible chromatin regions in our sorted cell types, we analyzed the ATAC-seq signal at every peak across all samples using the DE-Seq algorithm (Anders and Huber, 2010). From a total of 116,942 ATAC-seq peaks, we found 10,687 that have significant accessibility changes between samples (FDR <0.001). *k-*means clustering of differentially open peaks revealed three main groups of genomic regions that represent the open chromatin profiles of distinct cell states (**Figure 5A, Supplementary Table 7**). In Group I we observed 2,754 accessible regions in duct cells (either wild-type or Neurog3-null) that switch to a closed state in Neurog3^pos^ progenitors and remain closed in endocrine cells. Using the GREAT algorithm (McLean et al., 2010), we found that these regions were associated with genes that have established roles in exocrine pancreas cell development, gland development and cell proliferation like *Fgfr, Smad, Ptf1a, Hes1*, and Notch signaling (**Figure 5B**). Group II includes 6,312 and Group III includes 1,621 accessible regions (**Figure 5A**). Based on the ATAC-seq signal, we observed that these regions are closed in duct cells, open in Neurog3^pos^ progenitors and remain in open state in endocrine cells. The regions in Group III have significantly stronger ATAC-Seq signal in endocrine cells compared to endocrine progenitors, suggesting that other regulatory factors independent of Neurog3 might be enhancing the accessibility in these regions once the cells begin producing hormones. GREAT analysis linked chromatin from Groups II and III to genes known to regulate endocrine pancreas differentiation, or cardinal features of islet function including peptide hormone processing, and regulation of calcium ion-dependent exocytosis (**Figure 5B**).

**Figure 5:**
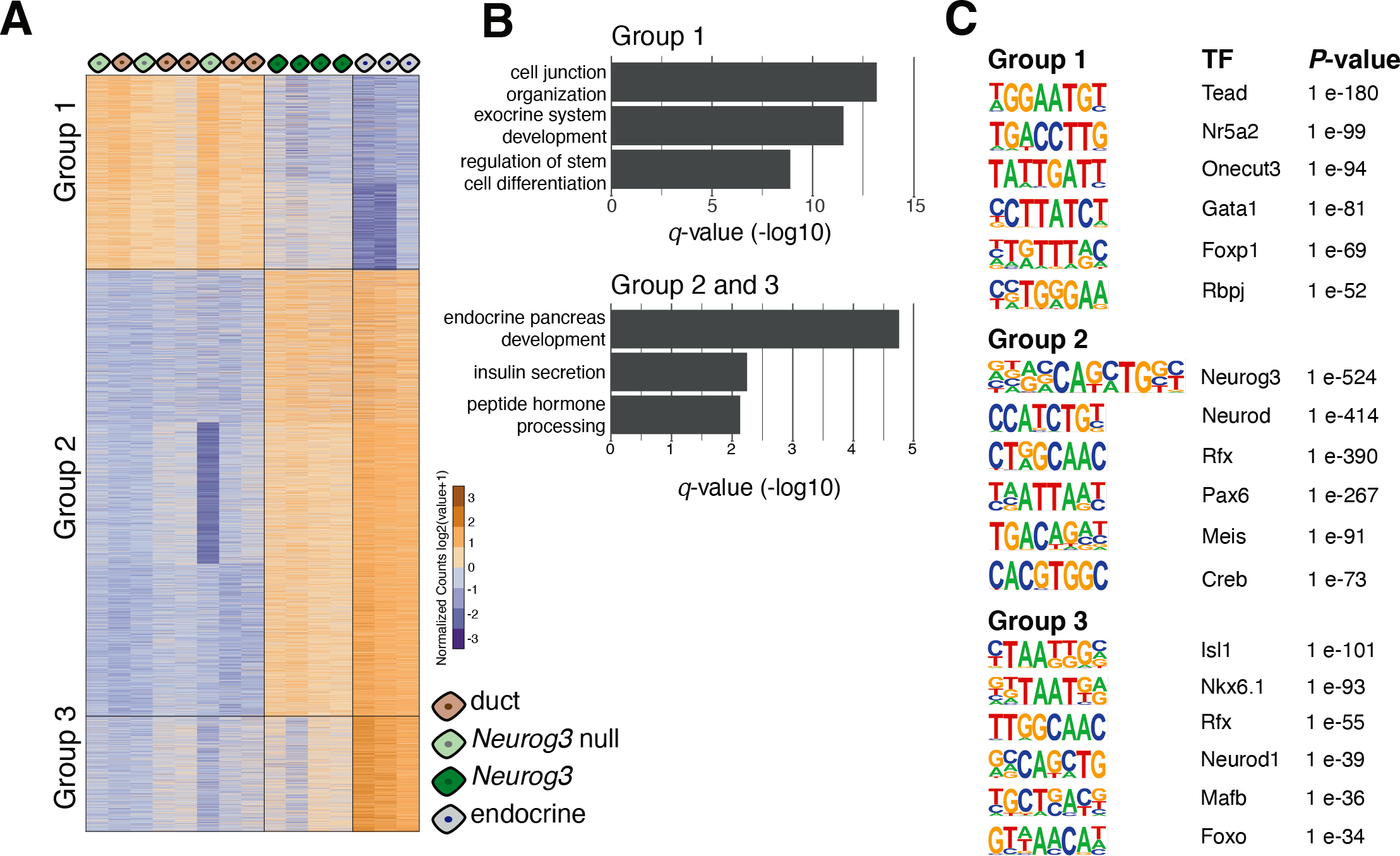
(A)Heat map shows differentially open chromatin regions. Each column is an ATAC-seq sample, each row is an open chromatin region, organized by k-means clustering. Three groups of open regions were identified and indicated on the graph. (B)Bar graphs show significant GO Terms associated with open regions identified in (A). (C)Position weight matrices of enriched TF motifs found in each of the three open chromatin groups.

To discover TF motifs within these dynamic chromatin regions, we performed TF motif enrichment analysis using the HOMER algorithm (Heinz et al., 2010). Consistent with GREAT analysis, we found overrepresented motifs (**Figure 5C**) of exocrine lineage specific factors like *Tead*, *Rbpj* and *Nr5a2* in accessible chromatin regions of duct cells in Group I. In contrast, our analysis of regions in Group II identified *Neurog3*, *NeuroD*, *Rfx* and *Pax* motifs— all known regulators of endocrine pancreas development. Likewise, the analysis of Group III regions yielded enriched TF motifs of lineage markers of *β*- and *α*-cells, including *Mafb* and *Isl1*. Thus, by combining cell sorting, mouse genetics, and ATAC-seq we identified developmentally resolved chromatin states, and found sequence motifs enriched for regulators of pancreas development, demonstrating the sensitivity and specificity of our approach.

### Identifying TF occupancy in regulatory genomic regions during endocrine cell differentiation

Chromatin accessibility assays, like ATAC-seq and Dnase-Seq, enable identification of TF occupancy sites where DNA is protected from enzymatic cleavage or transposition due to TF binding, leaving a “TF footprint” (Buenrostro et al., 2013; Maurano et al., 2012). We envisioned that an integrative approach combining TF footprint and single-cell gene expression profiles could uncover TF activity during endocrine pancreas differentiation. We used the BaGFoot algorithm to identify changes in TF occupancy between two cell states using our ATAC-seq samples (Baek et al., 2017). BaGFoot calculates two parameters for each TF motif: (1) footprint depth (**FPD**), the relative protection of DNA at the TF motif site, and (2) flanking accessibility (**FA**), the quantification of accessible chromatin near the TF motif (**Figure 6A**). TF binding dynamics is expected to affect these two parameters genome-wide; thus, by comparing the FPD and FA between two samples, we can infer changes in TF activity. For instance, a motif with a deep FPD, and high FA would indicate strong protection at the motif site. These results are represented in “bagplots”, which are analogous to “box and whisker” plots (**Figure 6B**, also see **Methods**).

**Figure 6:**
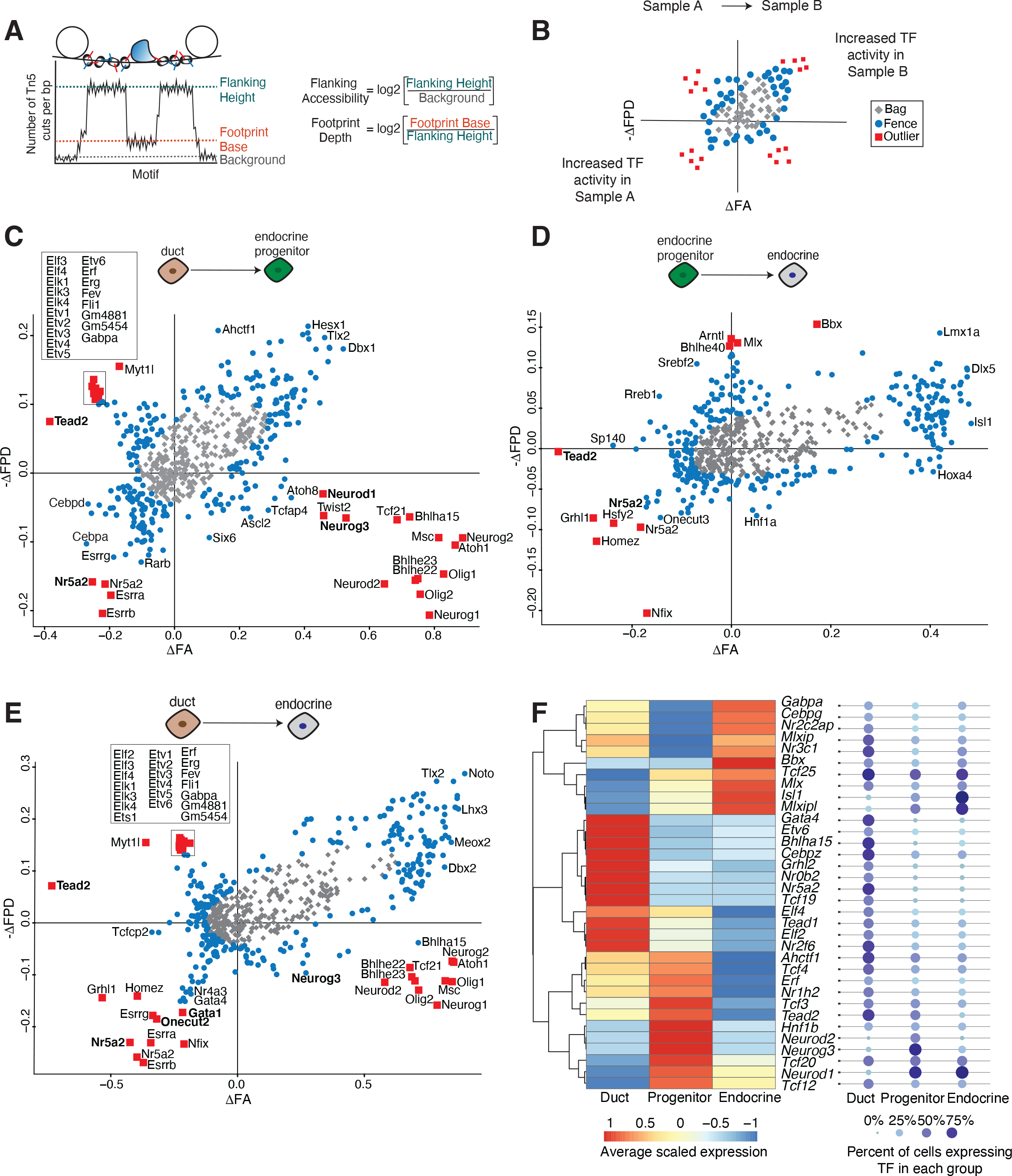
(A) Cartoon describing how footprint depth (FPD) and flanking accessibility (FA) are calculated from ATAC-seq data. (B) Guide to interpret pairwise comparisons using a bagplot. (C-E) Bagplots displaying TFs with upregulated activity when comparing two samples. Outliers are marked by red squares, TFs in the fence are marked by blue circles, and TFs in the bag are marked by grey diamonds. Bolded TFs correspond to the de novo motifs found in the HOMER analysis. (F) Heat map shows average expression levels of outlier TFs in duct, progenitor, or endocrine cells. TFs are ordered by hierarchical clustering, expression levels are scaled to each row. Each TF is detected in at least 25% of cells in each group.

We calculated the FPD and FA values for more than 650 curated TF motifs using our ATAC-seq data. Pairwise comparison of footprint signatures in duct cells and Neurog3^pos^ progenitors, or duct cells and endocrine cells revealed changes in TF activity. Consistent with the HOMER-based motif analysis, we found strong footprint signals for Gata and Onecut TFs, and Nuclear Receptors in duct cells. In endocrine cells, we detected footprints for homeobox TFs including *Isl1*, *Hnf1a* and *Pou* TFs (**Figure 6 C-E**, **Supplementary Table 8**). Comparison of Neurog3^pos^ progenitors and endocrine cells revealed relatively modest TF activity changes (**Figure 6D**).

Similar to the findings above, the most significant changes in TF footprint activity occurs during the transition from ductal to endocrine progenitor state, supporting the view that activation of Neurog3 is the main driver of changes in chromatin accessibility and gene expression.

We also calculated the FA and FPD scores of the TF motifs we derived *de novo* from our ATAC-seq motif enrichment analysis (**Figure 5C**). These motifs displayed increased FA or FPD in the appropriate cell type (indicated in bold, **Figures 6C-E** and **Supplementary Table 8**), independently validating the TF occupancy at these sequences.

While footprint depth and flanking accessibility are often correlated, some TFs only exhibited increased flanking accessibility without a detectable footprint, likely due to distinct DNA binding kinetics— for instance, those TFs with high OFF rates (Baek et al., 2017; Corces et al., 2018).

TFs matching this profile were basic helix-loop-helix (bHLH) factors including *Neurog3*, *Neurod1* and *Ascl2* in endocrine progenitors. In addition, some motifs were found in the second quadrant, displaying deeper FPD, but decreased FA in endocrine or Neurog3^pos^ progenitor samples compared to duct cells. This profile is consistent with repressor TFs, whose DNA binding activity leads to decreased accessibility surrounding the motif. We found that Tead factors and ETS family TFs, including *Etv6*, *Elf2/4*, *Erf* were included in this group (**Figure 6C, 6E**).

Paralogous TFs often bind similar DNA motifs, resulting in nearly identical footprint scores. For instance, *Neurog3* motif could also be recognized by *Neurog1* or *Neurog2* (**Figure 6C, E**). Thus, footprint analysis alone cannot determine which TF family member might be occupying the regulatory sequences in a particular cell type. Integrating BaGFoot results with single-cell expression data overcomes this limitation. We found more than 50 TFs whose expression correlates with a matching footprint (**Figure 6F, Supplementary Figure 5, Supplementary Table 9**). Among the TFs whose expression was detected in at least 25% of the cells within each group (**Figure 6F**), we confirmed the activity of known regulators, for instance *Nr5a2 and Gata4* in duct cells (Hale et al., 2014; Xuan et al., 2012). In addition, we found footprints of several relatively less-studied Nuclear Receptor TFs (*Nr2f6*, *Nr3c1*) and we identified a CTF/NFI factor, *Nfix* that has increased activity in Neurog3^pos^ progenitor cells (**Figure 6D, Supplementary Figure 5**). Taken together, footprint and expression analysis predicted dozens of regulators whose roles have not been previously explored in endocrine cell development, and provided quantitative evidence of selective TF occupancy in different pancreatic cell types.

## DISCUSSION

Here, we established an integrative approach combining cell purification, genetic labeling, single-cell transcriptomics, chromatin accessibility assessment and TF footprint analysis to elucidate molecular mechanisms underlying pancreatic endocrine cell specification. We show that endocrine cell development is a dynamic process involving a network of TFs whose expression is selectively tuned to define specific hormone lineages. We were able to delineate gene expression changes leading to *δ*-cell specification, and nominate unrecognized factors that could regulate *δ*-cell function. We demonstrate that in developing pancreatic epithelial cells, chromatin undergoes substantial reorganization upon Neurog3 induction. In remodeled genomic regions during development, we identified enriched TF motifs and footprints that correspond to TF activity in specific cell types.

A few prior studies (Scavuzzo et al., 2018; Yu et al., 2019) postulated that the *Neurog3^pos^* progenitors exhibit heterogeneity and temporal lineage biases. In our study, using the same mouse models and embryonic stages, we did not find evidence for such bias even though our gene expression results aligned well with differential gene expression reported by Scavuzzo and colleagues. Thus, differences in our findings may reflect interpretation of alternative analytical approaches, rather than primary data. Similar points about challenges in single-cell analysis and biological interpretation were discussed in recent reviews (Kiselev et al., 2019; Tritschler et al., 2019).

Using an iterative, semi-supervised clustering approach, we successfully identified branching points that specify three hormone lineages, including *β*-, *α*-, and *δ*-cell lineages. In our dataset we found only 13 PP cells, which did not provide sufficient statistical power to permit a PP-branch identification. Due to the known regulatory role of TFs, we focused on differentially expressed TFs between these lineages. We identified known, as well as previously under- studied pancreatic TFs that may have roles in islet endocrine cell specification. Based on TF expression in specific developmental timelines, we generated a network and observed that lineage specification is governed by a network of TFs with dynamic, overlapping expression profiles. For instance, while Neurog3 is necessary for the endocrine lineage, it needs to be turned off to permit further differentiation of endocrine cell lineages. We speculate that this may explain the low efficiency observed in direct reprogramming approaches when a handful of lineage-specific TFs are constitutively overexpressed to force non-islet cells toward a *β*-cell fate (Hickey et al., 2013; Li et al., 2014). Our focused analysis on *Neurog3^pos^* cells revealed that the pan-endocrine state precedes specific endocrine lineages, and the early endocrine cells are polyhormonal as defined by their transcriptome. This may explain why the interconversion of hormone cell types does not require Neurog3 (Chakravarthy et al., 2017; Furuyama et al., 2019). These results are also reminiscent of reports of polyhormonal cells generated during the in vitro differentiation experiments using human embryonic stem cells or adult tissues with endoderm origin (Galivo et al., 2017; Krentz et al., 2018; Lee et al., 2013; Petersen et al., 2017; Veres et al., 2019).

Chromatin accessibility is thought to be a better predictor of cell identity than transcriptome analysis, with changes in chromatin states often preceding changes in gene expression (Corces et al., 2016). By taking advantage of established cell markers and genetic models, we were able to dissect the chromatin accessibility changes during endocrine cell differentiation at unprecedented resolution. The unexpected similarity between duct cells and those that activate *Neurog3* forces a re-evalution of extant endocrine cell development models. For example, our findings provide evidence that pancreatic ‘trunk cells’ previously postulated to be oligopotent progenitors are simply duct cells that default to the ductal lineage in the absence of Neurog3 (**Figure 4E**). Comparison of *Neurog3^po^*^s^ cells from heterozygous (*Neurog3*^+/eGFP^) and homozygous wild-type (*Tg(eGFP);Neurog3^+/+^*) mice showed that a single, wild-type *Neurog3* allele is sufficient to drive global chromatin reconfiguration in the pancreatic endocrine lineage. It is likely that in individual ductal epithelium cells, Neurog3 concentration needs to reach a critical threshold to achieve pioneering activity and to compete with histone proteins for DNA binding (Bankaitis et al., 2015; Klemm et al., 2019).

Using a TF footprint algorithm, we provide quantitative, cell type-specific TF occupancy profiles at nucleotide resolution in pancreatic duct, endocrine progenitor and endocrine cell regulatory DNA. To our knowledge, this is the most comprehensive analysis of TF activity correlated with gene expression during pancreas development. TF-regulatory DNA interactions form the basis of gene regulatory networks, which are central to determining and maintaining cell type-specific transcription, cell fate and function. Further delineation of gene regulatory networks defining pancreatic cell lineages will be crucial for understanding pancreas disorders, and have the potential to improve gene therapy approaches using CRISPR-guided synthetic engineering to generate cells and tissues (Bevacqua et al., 2021). Expanding these strategies to human pancreas or in vitro differentiation efforts using emerging single-cell technologies that query chromatin and gene expression profiles (Ma et al., 2020) could offer new approaches to investigating the pathogenesis of type 1 and type 2 diabetes.

## METHODS

### Animal models

All animal experiments were conducted in accordance with Stanford University IACUC guidelines. *Neurog3^eGFP/+^* knock-in reporter mice were a kind gift from Dr. Klaus Kaestner (University of Pennsylvania, USA) (Lee et al., 2002) and were maintained on a CD1 background. *Neurog3-Cre* mice were obtained from Guoqiang Gu (Vanderbilt University, USA) and maintained on a mixed background of C57BL/6 and CD1 (Gu et al., 2002). *Rosa-mTmG* (Muzumdar et al., 2007) mice were obtained from the Jackson Laboratories and maintained on a mixed background of C57BL/6 and CD1. *Tg-eGFP; Neurog3^+/+^* transgenic mice were a kind gift from Drs. Guoqiang Gu and Douglas Melton (Gu et al., 2004). Timed matings were used to obtain mice at embryonic day (E) E15.5 and E17.5 for experiments; observation of a vaginal plug was considered E0.5 for embryonic staging purposes. Both male and female mice were used in all experiments.

### Tissue processing and FACS

Pancreata were dissected from E15.5 and E17.5 embryos and checked for GFP using a fluorescence dissecting microscope. GFP^pos^ pancreata were then digested with Tryp-LE express (ThermoFisher, 12605-010) for 5 minutes at 37℃, with regular pipet agitation to disrupt tissue. The digestion reaction was stopped by adding FACS buffer, which contains Ca^2+^ and Mg^2+^ free PBS supplemented with 2% Bovine serum albumin and 10 mM EGTA. The cell suspension was filtered to remove debris using a cell 70-micron cell strainer (BD Biosciences). Red blood cells were eliminated from dissociated cells using an RBC lysis buffer (BioLegend). Cells were then stained with Aqua live/dead viability dye (Thermo Fisher) to exclude dead cells during sorting. Cells were incubated with a blocking solution containing FACS buffer and goat IgG (Jackson Labs, 1:20 dilution) prior to staining with cell surface antibodies. After blocking, antibody staining was performed on ice for 30 minutes using the following antibodies: biotin mouse anti-CD133 (13A4, 1∶100; eBioscience), Streptavidin-APC (1∶200; eBioscience). We also used CD45-PE-Cy7 (eBioscience) to label and exclude leukocytes. We previously showed that CD133 labels Neurog3^pos^ endocrine progenitors and duct cells (Sugiyama et al., 2007). By contrast hormone^pos^ islet cells that no longer produce Neurog3 are CD133^neg^. After exclusion of CD45^pos^ cells, the following gating strategies defined pancreas cell subpopulations: GFP^pos^CD133^neg^ cells were considered ‘endocrine’, GFP^pos^CD133^pos^cells were ‘Neurog3^pos^’ or ‘Neurog3 null’ if obtained from null animals, and GFP^neg^CD133^pos^ cells were considered ‘duct’ (Benitez et al., 2014; Sugiyama et al., 2007). Representative gates are shown in Figure 4B. Note that the GFP intensity of *Neurog3*-null cells is reduced. In wild type cells, Neurog3 normally enhances its own expression through an auto-regulatory “positive feedback loop”. In null cells this mechanism is most likely absent (Ejarque et al., 2013; Lee et al., 2002; Wang et al., 2008).

### Single-cell RNA Sequencing

Single-cell RNA-seq libraries were generated using the SMART-Seq2 method as described (Picelli et al., 2014). Dissociated cells were sorted directly into 96-well plates containing lysis buffer with ERCC RNA spike-in controls (ThermoFisher). The details about the sorted cell populations, genotypes, and associated plate codes are available in the GEO metadata file linked to this study. The lysis reaction was followed by reverse transcription with template-switch using an LNA-modified template switch oligos to generate cDNA. After pre-amplification, DNA was purified and analyzed on an automated Fragment Analyzer (Advanced Analytical). cDNA fragment profile corresponding to each single cell was individually inspected and only wells with successful amplification products (concentration higher than 0.06 ng/ul) and no detectable RNA degradation were selected for final library preparation. Tagmentation assays and barcoded sequencing libraries were prepared using Nextera XT kit (Illumina) according to the manufacturer’s instructions. Barcoded libraries were pooled and subjected to 75 bp paired-end sequencing on the Illumina NextSeq instrument.

### RNA-Seq Read Alignment

Raw reads passing quality control using FastQC were aligned to a custom reference genome consisting of Fasta files for mm10, ERCC spike in controls, and three transgenes: *eGFP*, *tdTomato*, and *Cre*. STAR was used to create the custom genome and read alignment (Dobin et al., 2012). The resulting BAM/SAM files were used to create a ‘master counts table’ using HT- seq (Supplementary Table 1) (Anders et al., 2015). Cells had an average of 3,044 genes expressed per cell, ranging from 1,237 to 6,047 genes.

### Unsupervised Single-cell Clustering and Trajectory Analyses

Clustering and trajectory analysis were performed using the single-cell analysis package Monocle 2 (v. 2.4.0) (Qiu et al., 2017b). A flowchart summarizing each analysis step is provided in Supplementary Figure 2. Before starting the analysis, the transgenes *GFP, Cre* and *Td- tomato* were removed from the master counts table (Supplementary Table 1). Unsupervised clustering aims to cluster the cells based on global gene expression profiles. First step is to choose which genes to use to cluster the cells. Based on the dispersion calculations, we set the mean_expression parameter to 1. Before performing dimension reduction, the data was examined using the `plot_pc_variance_explained` function, which plots the percentage of variance explained by each principal component on the normalized expression data. Based on the ‘elbow’ method, we determined that the first 5 dimensions showed the majority of data variability. Therefore, *t*-distributed stochastic neighbor embedding (*t*-SNE) dimension reduction was performed on the first 5 principal components. We set *num_clusters* to 7 to visualize cell clusters (Supplementary Figure 1). The identity of the cell clusters was revealed by mapping marker gene expression levels onto single-cells (Figure 1A-B). Clusters 4 and 6 were combined and labeled “Endocrine 1”. At this point, the 14 mesenchymal cells that formed Cluster 5, and genes that were expressed in less than 5 cells were filtered out. To establish pseudotime trajectories, Monocle’s `differentialGeneTest` function was used to find genes that vary among the clusters, specified as `fullModelFormulaStr = “∼Cluster”.` Top 100 genes with the lowest *q*-value were used to order cells, and a pseudotime trajectory was constructed using the DDRTree method. To identify gene expression changes between cells aligned along the established pseudotime trajectories, we used Monocle’s `differentialGeneTest` function by specifying `fullModelFormulaStr = “∼Pseudotime”.` We considered genes significant if the rounded *q*-value was less than or equal to 0.05. Gene ontology terms were found for each of the 7 clusters using DAVID v6.8 (Supplementary Table 3) (Huang et al., 2009).

### Semi-supervised Single-cell Clustering and Trajectory Analyses

Semi-supervised clustering and trajectory analyses were performed to resolve individual endocrine lineage branching (Supplementary Figure 2). The process begins with defining marker genes that represent cell populations, then identifying the genes that co-vary with these markers, and finally ordering the cells based on these co-varying genes. Monocle provides the `CellTypeHierarchy` function for semi-supervised clustering analysis. Since our goal was to resolve the *β*-, *α*- and *δ*-cell branches, we picked marker genes as *Neurog3* for endocrine progenitors, *Ins1* and *Ins2* for *β*-cells, *Gcg* for *α*-cells, and *Sst* for *δ*-cells. We set the expression threshold in each cell for these markers to 100 or more reads. Accordingly, cells that express more than one marker gene are labeled “ambiguous” and cells that do not fit into any marker gene category are labeled as “unknown”. The gene list was further filtered to remove genes if detected in less than 5 cells. Top 100 genes that co-varied with the marker genes (400 genes in total) were considered for the clustering and trajectory analysis. Note that the semi-supervised analysis was limited to the 317 cells that were placed after the *Neurog3* peak expression in the unsupervised trajectory, which corresponds to the pseudotime point 6.7. The first iteration separated *β*-cells in one branch and the majority of *α*- and *δ*-cells in a second branch. To split the *α*- and *δ*-branches, we again focused on cells of interest, and excluded the cells on the *β*- cell branch to create a new `CellDataSet (cds)` object in Monocle. In this new `cds()` object, cells were relabeled as *α*-, *δ*-, and *Neurog3^pos^* cells based on marker gene expression.

Trajectory analysis was performed as described earlier. The final iteration established trajectories with *α*- and *δ*-cells separated on own branches. Similar to unsupervised clustering, Monocle’s `differentialGeneTest` (by specifying `fullModelFormulaStr = “∼Pseudotime”`) function was used to identify genes whose expression changes significantly during each endocrine lineage specification. For differential gene expression analysis, cells with pseudotime point > 5.7 and ≤ 6.7 were also included (peak *Neurog3* expression) to visualize the cell fate transitions beginning from the *Neurog3^pos^* progenitors. Hence, three differential gene tests were performed to determine transcriptome changes from *Neurog3*^pos^ progenitor cells to each of the three endocrine lineages. Results from differential expression analyses were filtered to include genes with a *q*-value less than 0.1 and those in the top 50% of normalized base mean expression among cells within each branch. All differentially expressed genes lists were further narrowed to TFs for a total of 145 TFs. These TFs are visualized in a heatmap where all cells were aligned in pseudotime order (Supplementary Figure 4).

### Analysis and classification of *Neurog3^pos^* progenitors

The master read counts table (Supplementary Table 1) was subset to select *Neurog3^pos^* cells. We defined *Neurog3^pos^* cells as any cell with at least 10 read counts for *Neurog3*, resulting in 214 cells. The semi-supervised clustering approach was used to label and cluster cells based on either *Neurog3* or *Chga* expression (see the previous section). Top 100 genes that co-varied with the marker genes (200 genes in total) were considered for the clustering and trajectory analysis. *t*-SNE dimension reduction was performed on the first two principal components, and *num_clusters* was set to 3. Based on the *Neurog3* levels, the clusters were named High, Medium, and Low. A trajectory was established by finding differentially expressed genes among the High, Medium, Low clusters, using Monocle’s `differentialGeneTest` function by specifying `fullModelFormulaStr = “∼Cluster”`. Top 100 genes with the lowest *q*-value were used to order cells, and a pseudotime trajectory was constructed using the DDRTree method. The trajectory was colored based on embryonic day (Figure 2F) or cluster (Figure 2H).

To count hormone expressing cells, we analyzed the read counts of *Ins1*, *Ins2*, *Gcg* and *Sst* in each *Neurog3^pos^* cell. Any detectable expression (i.e. size-factor normalized counts > 0) was counted. The cells were then categorized as expressing zero, one, two or three hormones (*Ins1* and *Ins2* reads were combined and presented as *Ins*).

### Expression Specificity Scores, TF-Cell Type/State Network

We derived expression specificity scores for TFs that are differentially expressed during endocrine cell lineage specification. We have previously used this method to reveal cell type-specific gene expression in human pancreas cells (Arda et al., 2018). ESS was calculated as follows:

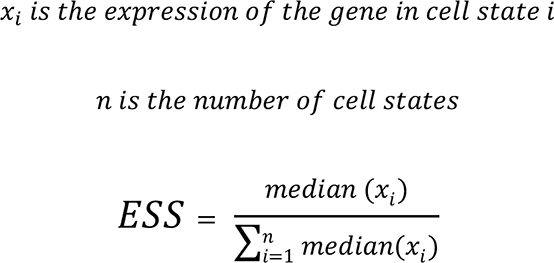

A cell state is defined here as population of cells that are quantitatively distinct based on their transcriptome. Two cell states (early progenitor, late progenitor) and four cell types (duct, *β*-, *α*- and *δ*-cells) were used to determine the expression specificity score of each TF. The duct cells were categorized as cells with pseudotime values < 3 (53 cells) based on the unsupervised trajectory analysis. Early progenitor state has cells with pseudotime values between 3 and 6.7 (111 cells). Late progenitor state cells have a pseudotime value greater than 6.7 and include those that were not assigned to an endocrine lineage (121 cells). The hormone producing cells consist of those assigned to their respective endocrine cell branch (90 cells in the *β*-lineage, 76 cells in the *α*-lineage, and 30 cells in the *δ*-lineage). To obtain *x*_*i*_, we used the size-factor normalized single-cell RNA-Seq counts as gene expression values. Thus, a TF with an ESS of zero would indicate no expression in that cell type/state, and an ESS of 1 would indicate exclusive expression, i. e. the TF is only expressed in that cell state. We obtained the list of differentially expressed TFs by overlapping the gene the list with a curated TF list (Weirauch et al., 2014), yielding 145 TFs. The TF list was further narrowed to 87 by only including those that were detected in at least 50% of the cells in that cell type/state (Supplementary Table 5). The network was generated by Cytoscape (version: 3.8.2) (Shannon et al., 2003). The color and thickness of the network edges (connections) directly corresponds with the expression specificity score (ESS) of the TF in the interacting cell type/state.

### ATAC-seq assays and data processing

Three mouse genotypes were used for ATAC-seq analysis, *Tg-eGFP; Neurog3^+/+^*, *Neurog3^eGFP/+^*, and *Neurog3^eGFP/eGFP^*. From these animals, different cell populations were isolated as described in the ‘Tissue processing and FACS’ section (also see Supplementary Table 6). ATAC-seq was performed following the protocol in Buenrostro et al., 2013. On average 10,000 sorted cells were used for each ATAC-seq assay. Sorted cells were pelleted at 300 *g*, washed once with PBS. Nuclei were isolated, followed by the transposition reaction.

Transposed DNA fragments were purified using the Qiagen MinElute kit and amplified 6-8 cycles using the Nextera (Illumina) PCR primers. Libraries were sequenced as 2x50 on HiSeq2000 platform. ATAC-seq data processing and genome alignment was performed with PEPATAC (version 0.8.2), a pipeline developed to analyze ATAC-seq samples (Smith et al., 2021). PEPATAC begins by trimming adapters using skewer (version 0.2.2) with the parameters “-f sanger -t 8 -m pe”. Trimmed fastq files were then mapped to the mm10 genome with bowtie2 (Langmead et al., 2009) and the parameter “--very-sensitive”. Lastly peaks were called using MACS2 (Feng et al., 2012) with “-q 0.01 --shift 0 –nomodel”. At the end of PEPATAC processing, 42-88 million reads aligned to the mouse genome and 15,377-55,676 peaks per sample were detected. These peak regions were then merged using BedTools (Quinlan and Hall, 2010) to generate a non-overlapping consensus peak list for downstream analysis. ATAC-seq fragments corresponding to the peaks were quantified by using the `annotatePeaks.pl` function in HOMER suite, a genome analysis tool (v.4.10) (Heinz et al., 2010). DE-Seq (Anders and Huber, 2010) was used to find regions with significantly different ATAC-seq counts by running a generalized linear model with the modelFormula set to “count∼condition” and “count∼1”. Accordingly, DE-Seq calculates *P*-values and FDR. Peaks passing the FDR threshold < 0.001 were considered ‘differentially open regions’ (DORs) between cell types (∼10,600 DORs). Pearson correlation coefficient method was used to determine the similarity between ATAC-seq samples based on DORs. The results were visualized using the R package `ggcorrplot` with hierarchical clustering. DORs and samples were clustered by Cluster 3.0 tool using the *k*-means method (de Hoon et al., 2004). ATAC-seq fragment counts were further normalized by log2 transformation after shifting values +1 for visualization in TreeView (Saldanha, 2004). To assign DORs to regulatory domains and putative target genes we used the GREAT algorithm (v3.0.0) (McLean et al., 2010) with default settings. GREAT also outputs enriched GO Terms associated with these regions. For the GO Term enrichment analysis, DORs were used as test regions against whole genome (mm10) as background.

### TF motif enrichment analysis

HOMER’s `findMotifsGenome.pl` function with ‘size 500 -len 6,8’ options was used to find enriched TF motifs in each DOR group (Heinz et al., 2010). HOMER’s *de novo* motif discovery analysis outputs a position weight matrix (PWM) for each significant motif. These PWMs were queried in the CisBP database (Weirauch et al., 2014) to find transcription factors associated with the significant motifs.

### BaGFoot analysis and integration of gene expression

BaGFoot footprint analysis was performed as described in (Baek et al., 2017). Narrow peaks were called for all ATAC-seq samples using MACS2 (Feng et al., 2012) and merged to generate a set of consensus peaks for BaGFoot. Peaks overlapping with black listed regions (downloaded from http://mitra.stanford.edu/kundaje/akundaje/release/blacklists/mm10-mouse/mm10.blacklist.bed.gz, Amemiya et al., 2019) were removed from the analysis. 662 mouse TF motifs were curated from TRANSFAC (Matys et al., 2006), JASPAR (Mathelier et al., 2016) and UniPROBE (Newburger and Bulyk, 2009). In addition, we included 19 *de novo* motifs derived from our ATAC-seq data by HOMER motif analysis. ATAC-seq sample replicates were grouped as follows: the duct dataset consisted of duct-het, duct-null, and Neurog3-null samples, the Neurog3 dataset consisted of Neurog3-het and Neurog3-Tg samples, and the endocrine dataset consisted of Endo-het and Endo-Tg samples. Each group were compared pairwise to detect TF footprint activity at motif locations. BaGFoot results are presented in “bag plots”, where each data point represents a TF motif. In a bag plot, the bag area contains 50% of the data (similar to the box in the box plot), the fence contains 97%-100% of the data points (similar to the whiskers in a box plot) (Rousseeuw et al., 1999). Any data point outside the fence is an outlier. Most TF motifs are not expected to be different between two conditions, and thus are localized around the origin. The significant motifs were statistically determined by Hotelling’s T-squared test and were labeled as outliers.

Based on the BaGFoot results, we compiled a list of outlier TFs (and their paralogs) to analyze their expression levels in the scRNA-Seq data. 481 cells were divided into duct, progenitor, and endocrine cell types to obtain average expression levels for outlier TFs. Cells were assigned to one of these three cell types based on their placement from the pseudotime trajectory analyses. Endocrine cells are a combination of cells aligned on the *β*-, *α*-, and *δ*- branch (Supplementary Table 9). The TFs whose expression was detected in at least 25 cells within each cell group were listed in Figure 6F. Those detected in fewer than 25% of the cells were shown in Supplementary Figure 5.

## ACKNOWLEDGMENTS

We thank P. Batista, the members of the Arda and Kim laboratories for discussions and suggestions. This work was supported by the Intramural Research Program of the NIH, National Cancer Institute, Center for Cancer Research, USA, by funds from a JDRF Advanced Postdoctoral Fellowship (3-APF-2016-172-A-N) to H.E.A., and by the National Institute of General Medical Sciences (R35-GM128636) to N.C.S. Computational resources of the NIH HPC Biowulf cluster supported the analysis in this work (http://hpc.nih.gov).

## Author contributions

H.E.A. and S.K.K. conceived and coordinated the study. C.M.B., P.T.P., L.L., K.T. bred the mice, performed embryonic pancreas dissection. H.E.A performed FACS, ATAC-seq, sc-RNA-Seq experiments. M.E., S.R.K, assisted with scRNA-Seq experiments and analysis. J.P.S. and N.C.S wrote the PEPATAC pipeline. S.B. wrote and performed the BagFoot analysis. E.D, H.E.A. analyzed the results, performed computational analysis. E.D., S.K.K. and H.E.A. interpreted the findings and wrote the manuscript with input from the other co-authors.

---

**Supplementary Figure 1.**
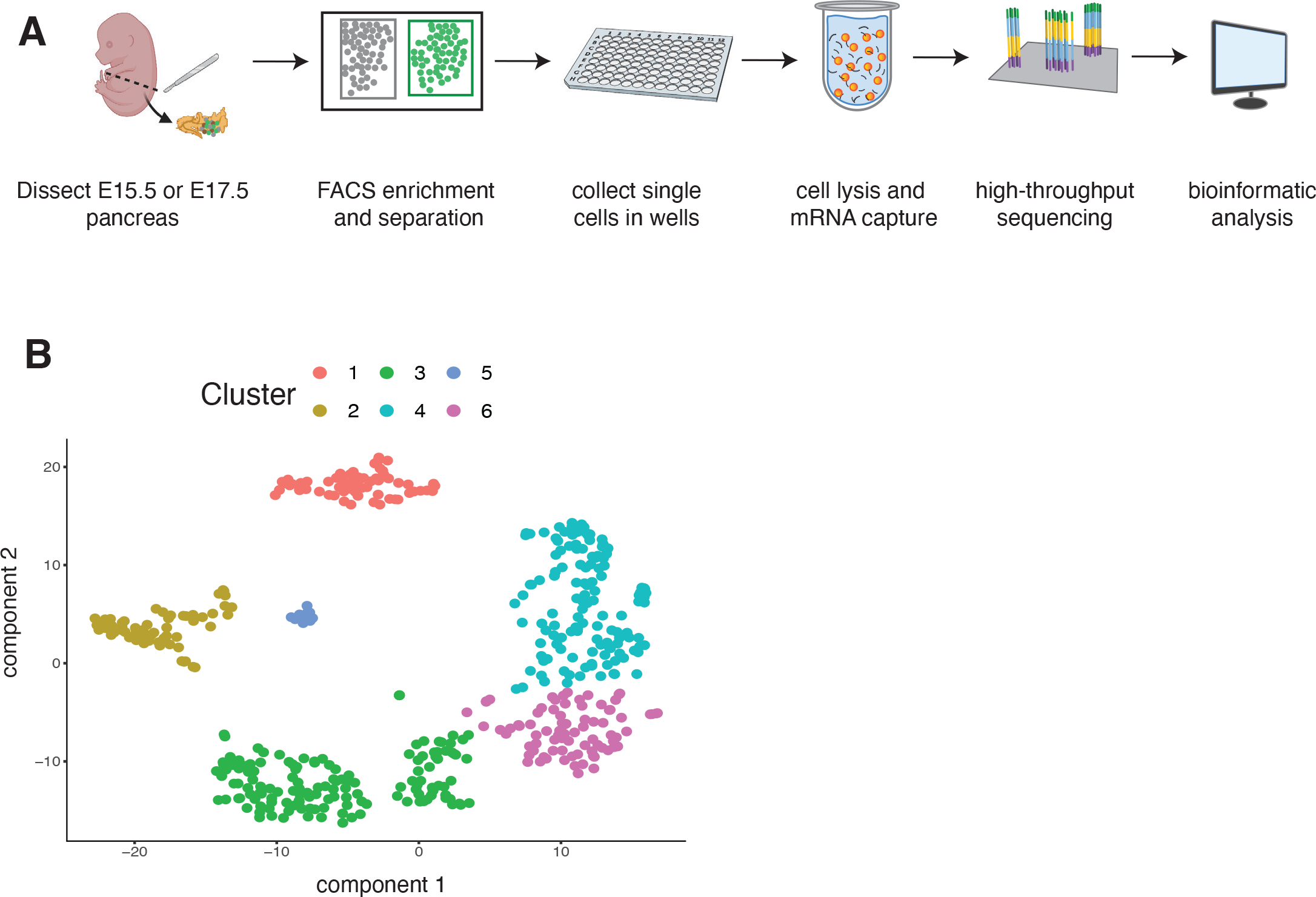
A) Illustration of the scRNA-Seq workflow performed in this study. See methods for details. B) t-SNE plot showing single cell clusters after unsupervised clustering approach.

**Supplementary Figure 2:**
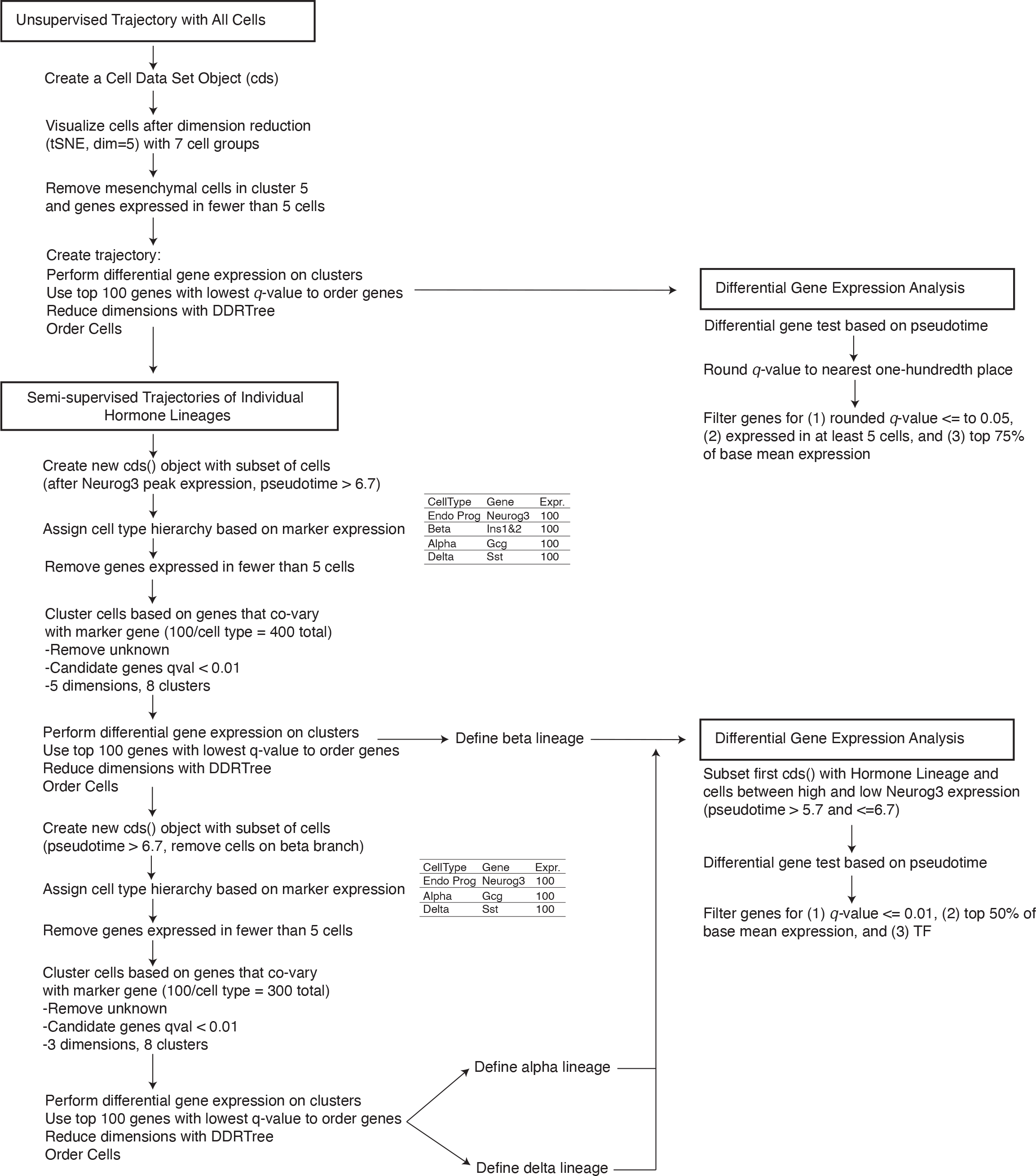
Flowchart summarizing the sc-RNA-Seq data analysis performed in this study using Monocle.

**Supplementary Figure 3.**
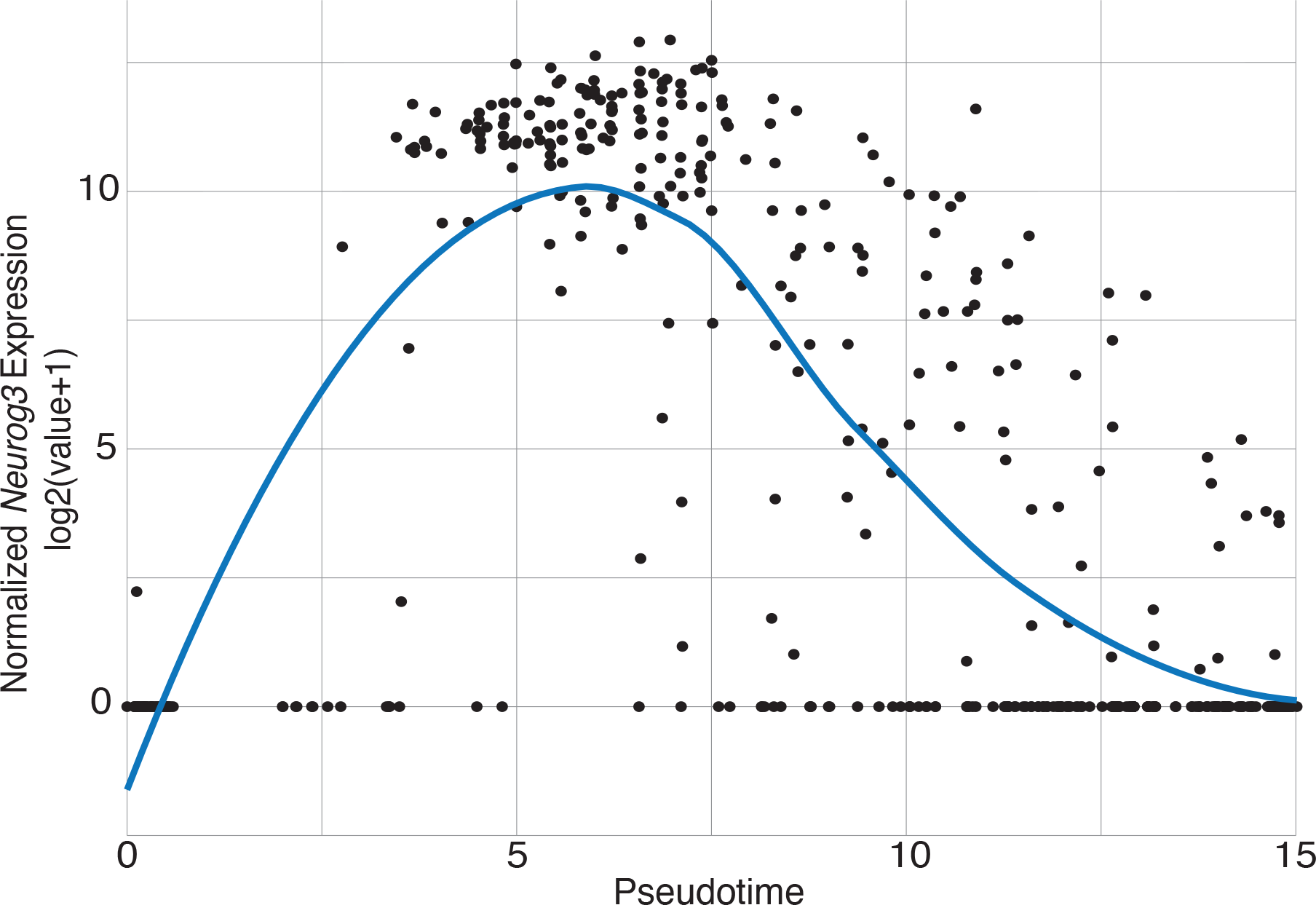
Distribution of Neurog3 transcript levels in single cells, ordered by the pseudotime defined in Figure 1C.

**Supplementary Figure 4.**
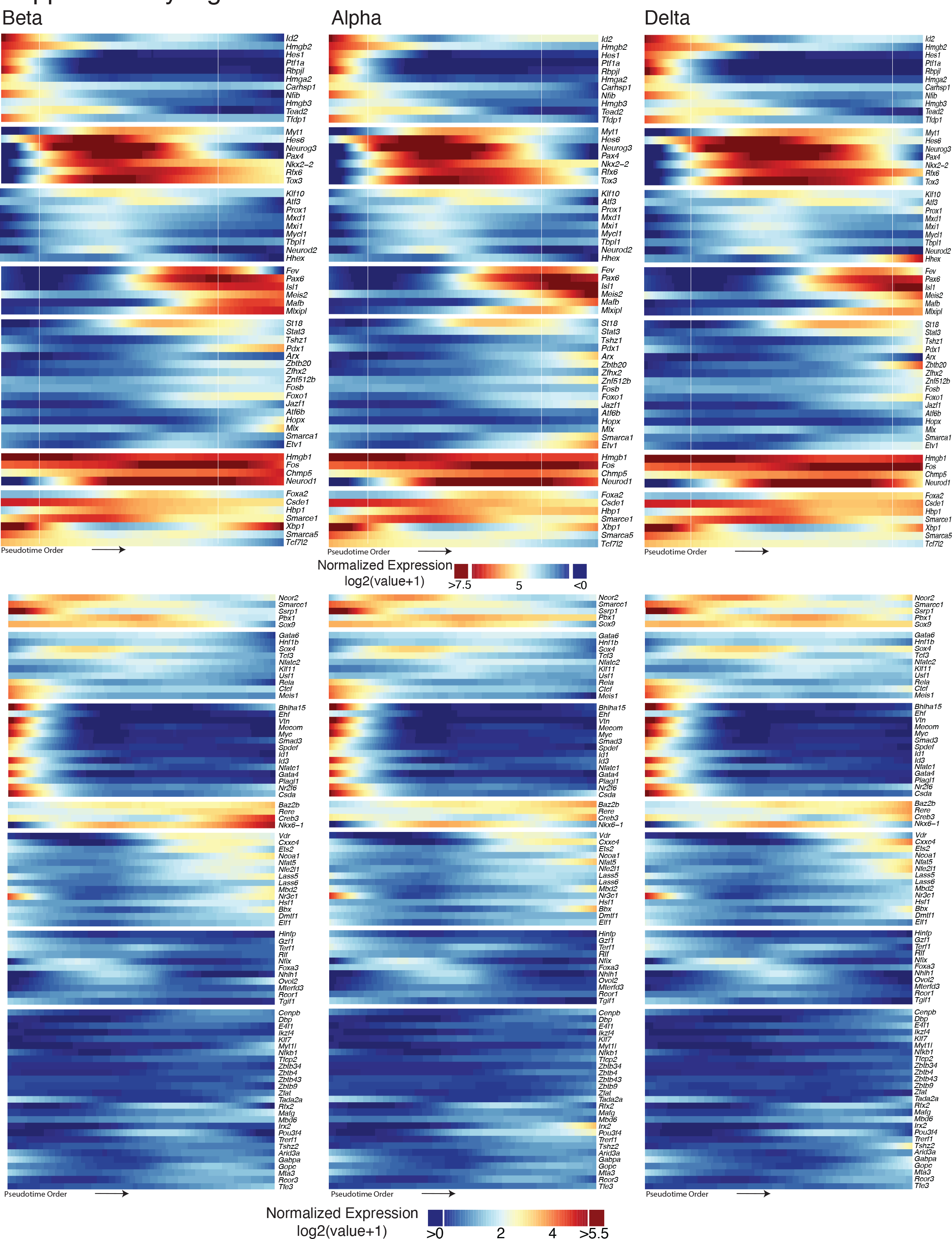
Heat maps showing dynamic expression level changes through endocrine cell differentiation beginning with *Neurog3^pos^* progenitor cells in *β*-, *α*-, or *δ*-branch. 145 TFs were found to be differentially expressed along this pseudotime order obtained from semi-supervised clustering analysis. For visualization purposes, the TF list is split into those with high (top) or low (bottom) expression. Color scale indicates log2 transformed normalized expression values after loess smoothing.

**Supplementary Figure 5.**
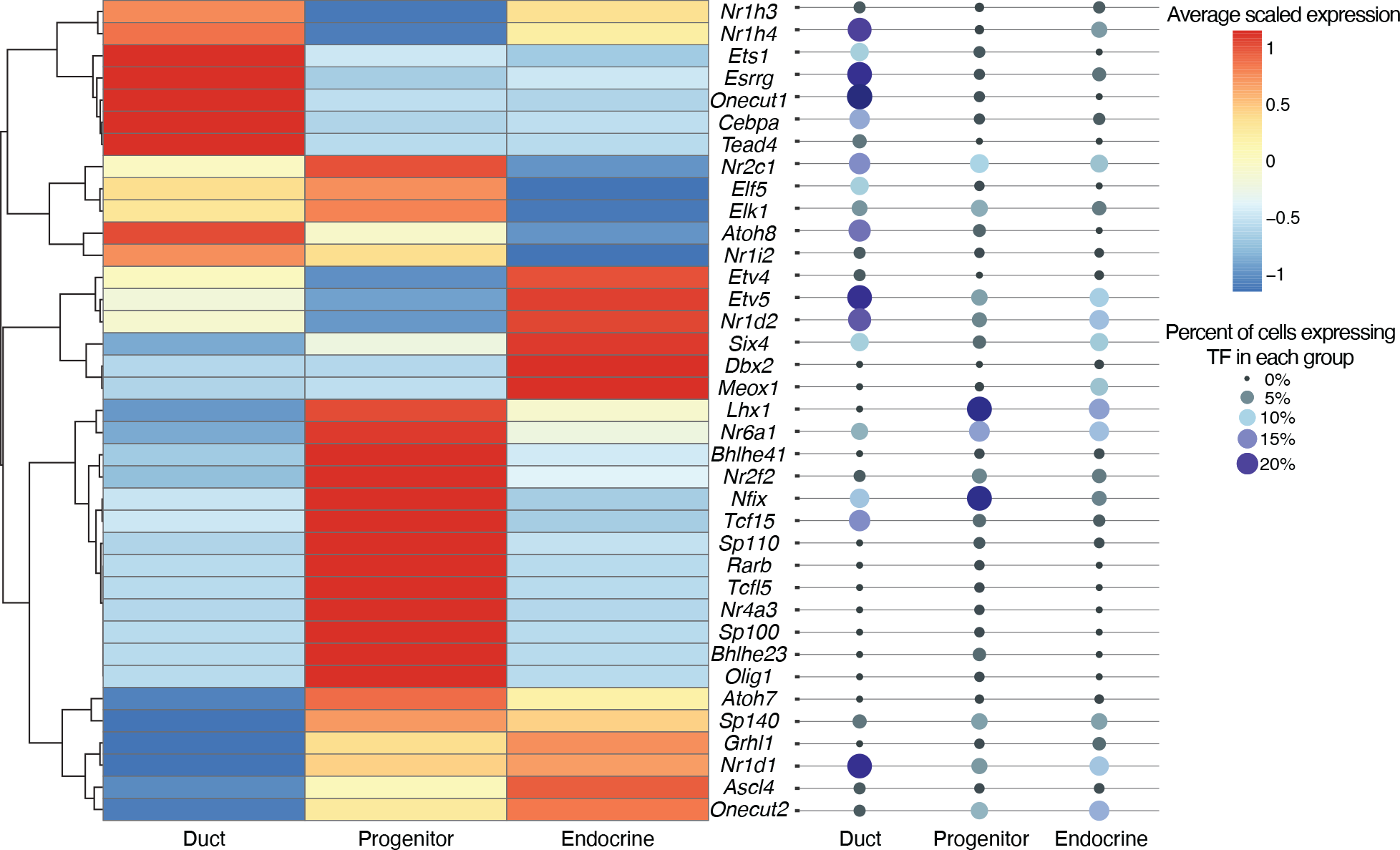
Heat map shows average expression levels of outlier TFs in duct, progenitor, or endocrine cells. TFs are ordered by hierarchical clustering; expression levels are scaled to each row. Each TF is detected in less than 25% of cells for each group.

## REFERENCES

1. Amemiya, H.M., Kundaje, A., and Boyle, A.P. (2019). The ENCODE Blacklist: Identification of Problematic Regions of the Genome. Sci Rep 9, 9354.

2. Anders, S., and Huber, W. (2010). Differential expression analysis for sequence count data. Genome Biology 11, R106.

3. Anders, S., Pyl, P.T., and Huber, W. (2015). HTSeq--a Python framework to work with high-throughput sequencing data. Bioinformatics 31, 166–169.

4. Arda, H.E., Benitez, C.M., and Kim, S.K. (2013). Gene regulatory networks governing pancreas development. Developmental Cell 25, 5–13.

5. Arda, H.E., Tsai, J., Rosli, Y.R., Giresi, P., Bottino, R., Greenleaf, W.J., Chang, H.Y., and Kim, S.K. (2018). A chromatin basis for cell lineage and disease risk in the human pancreas. Cell Systems 7, 310–322.

6. Arrojo e Drigo, R., Jacob, S., García-Prieto, C.F., Zheng, X., Fukuda, M., Nhu, H.T.T., Stelmashenko, O., Peçanha, F.L.M., Rodriguez-Diaz, R., Bushong, E., et al. (2019). Structural basis for delta cell paracrine regulation in pancreatic islets. Nat Commun 10, 3700.

7. Baek, S., Goldstein, I., and Hager, G.L. (2017). Bivariate Genomic Footprinting Detects Changes in Transcription Factor Activity. Cell Rep 19, 1710–1722.

8. Bankaitis, E.D., Bechard, M.E., and Wright, C.V.E. (2015). Feedback control of growth, differentiation, and morphogenesis of pancreatic endocrine progenitors in an epithelial plexus niche. Genes Dev. 29, 2203–2216.

9. Bastidas-Ponce, A., Scheibner, K., Lickert, H., and Bakhti, M. (2017). Cellular and molecular mechanisms coordinating pancreas development. Development 144, 2873–2888.

10. Bastidas-Ponce, A., Tritschler, S., Dony, L., Scheibner, K., Tarquis-Medina, M., Salinno, C., Schirge, S., Burtscher, I., Böttcher, A., Theis, F., et al. (2019). Massive single-cell mRNA profiling reveals a detailed roadmap for pancreatic endocrinogenesis. Development dev.173849.

11. Benitez, C.M., Goodyer, W.R., and Kim, S.K. (2012). Deconstructing pancreas developmental biology. Cold Spring Harb Perspect Biol 4.

12. Benitez, C.M., Qu, K., Sugiyama, T., Pauerstein, P.T., Liu, Y., Tsai, J., Gu, X., Ghodasara, A., Arda, H.E., and Zhang, J. (2014). An Integrated Cell Purification and Genomics Strategy Reveals Multiple Regulators of Pancreas Development. PLOS Genetics 10, e1004645.

13. Bevacqua, R.J., Dai, X., Lam, J.Y., Gu, X., Friedlander, M.S.H., Tellez, K., Miguel-Escalada, I., Bonàs-Guarch, S., Atla, G., Zhao, W., et al. (2021). CRISPR-based genome editing in primary human pancreatic islet cells. Nat Commun 12, 2397.

14. Buenrostro, J.D., Giresi, P.G., Zaba, L.C., Chang, H.Y., and Greenleaf, W.J. (2013). Transposition of native chromatin for fast and sensitive epigenomic profiling of open chromatin, DNA-binding proteins and nucleosome position. Nat. Methods 10, 1213–1218.

15. Byrnes, L.E., Wong, D.M., Subramaniam, M., Meyer, N.P., Gilchrist, C.L., Knox, S.M., Tward, A.D., Ye, C.J., and Sneddon, J.B. (2018). Lineage dynamics of murine pancreatic development at single-cell resolution. Nature Communications 9, 1–17.

16. Chakravarthy, H., Gu, X., Enge, M., Dai, X., Wang, Y., Damond, N., Downie, C., Liu, K., Wang, J., Xing, Y., et al. (2017). Converting Adult Pancreatic Islet α Cells into β Cells by Targeting Both Dnmt1 and Arx. Cell Metabolism 25, 622–634.

17. Corces, M.R., Buenrostro, J.D., Wu, B., Greenside, P.G., Chan, S.M., Koenig, J.L., Snyder, M.P., Pritchard, J.K., Kundaje, A., Greenleaf, W.J., et al. (2016). Lineage-specific and single-cell chromatin accessibility charts human hematopoiesis and leukemia evolution. Nature Genetics 48, 1193–1203.

18. Corces, M.R., Granja, J.M., Shams, S., Louie, B.H., Seoane, J.A., Zhou, W., Silva, T.C., Groeneveld, C., Wong, C.K., Cho, S.W., et al. (2018). The chromatin accessibility landscape of primary human cancers. Science 362, eaav1898.

19. Dobin, A., Davis, C.A., Schlesinger, F., Drenkow, J., Zaleski, C., Jha, S., Batut, P., Chaisson, M., and Gingeras, T.R. (2012). STAR: ultrafast universal RNA-seq aligner. Bioinformatics bts635.

20. Ejarque, M., Cervantes, S., Pujadas, G., Tutusaus, A., Sanchez, L., and Gasa, R. (2013). Neurogenin3 Cooperates with Foxa2 to Autoactivate Its Own Expression. Journal of Biological Chemistry 288, 11705–11717.

21. Feng, J., Liu, T., Qin, B., Zhang, Y., and Liu, X.S. (2012). Identifying ChIP-seq enrichment using MACS. Nature Protocols 7, 1728–1740.

22. Furuyama, K., Chera, S., van Gurp, L., Oropeza, D., Ghila, L., Damond, N., Vethe, H., Paulo, J.A., Joosten, A.M., Berney, T., et al. (2019). Diabetes relief in mice by glucose-sensing insulin-secreting human α-cells. Nature 567, 43–48.

23. Galivo, F., Benedetti, E., Wang, Y., Pelz, C., Schug, J., Kaestner, K.H., and Grompe, M. (2017). Reprogramming human gallbladder cells into insulin-producing β-like cells. PLoS ONE 12, e0181812.

24. Gradwohl, G., Dierich, A., LeMeur, M., and Guillemot, F. (2000). neurogenin3 is required for the development of the four endocrine cell lineages of the pancreas. Proc Natl Acad Sci U S A 97, 1607–1611.

25. Gu, G., Dubauskaite, J., and Melton, D.A. (2002). Direct evidence for the pancreatic lineage: NGN3+ cells are islet progenitors and are distinct from duct progenitors. Development 129, 2447–2457.

26. Gu, G., Wells, J.M., Dombkowski, D., Preffer, F., Aronow, B., and Melton, D.A. (2004). Global expression analysis of gene regulatory pathways during endocrine pancreatic. Development 131, 165–179.

27. Hale, M.A., Swift, G.H., Hoang, C.Q., Deering, T.G., Masui, T., Lee, Y.-K., Xue, J., and MacDonald, R.J. (2014). The nuclear hormone receptor family member NR5A2 controls aspects of multipotent progenitor cell formation and acinar differentiation during pancreatic organogenesis. Development 141, 3123–3133.

28. Heinz, S., Benner, C., Spann, N., Bertolino, E., Lin, Y.C., Laslo, P., Cheng, J.X., Murre, C., Singh, H., and Glass, C.K. (2010). Simple combinations of lineage-determining transcription factors prime cis-regulatory elements required for macrophage and B cell identities. Mol. Cell 38, 576–589.

29. Hess, D.A., Humphrey, S.E., Ishibashi, J., Damsz, B., Lee, A.-H., Glimcher, L.H., and Konieczny, S.F. (2011). Extensive pancreas regeneration following acinar-specific disruption of Xbp1 in mice. Gastroenterology 141, 1463–1472.

30. Hetz, C. (2012). The unfolded protein response: controlling cell fate decisions under ER stress and beyond. Nat Rev Mol Cell Biol 13, 89–102.

31. Hickey, R.D., Galivo, F., Schug, J., Brehm, M.A., Haft, A., Wang, Y., Benedetti, E., Gu, G., Magnuson, M.A., Shultz, L.D., et al. (2013). Generation of islet-like cells from mouse gall bladder by direct ex vivo reprogramming. Stem Cell Research 11, 503–515.

32. de Hoon, M.J.L., Imoto, S., Nolan, J., and Miyano, S. (2004). Open source clustering software. Bioinformatics 20, 1453–1454.

33. Huang, D.W., Sherman, B.T., and Lempicki, R.A. (2009). Systematic and integrative analysis of large gene lists using DAVID bioinformatics resources. Nat Protoc 4, 44–57.

34. Kiselev, V.Y., Andrews, T.S., and Hemberg, M. (2019). Challenges in unsupervised clustering of single-cell RNA-seq data. Nat Rev Genet 20, 273–282.

35. Klemm, S.L., Shipony, Z., and Greenleaf, W.J. (2019). Chromatin accessibility and the regulatory epigenome. Nat Rev Genet 20, 207–220.

36. Kopinke, D., Brailsford, M., Shea, J.E., Leavitt, R., Scaife, C.L., and Murtaugh, L.C. (2011). Lineage tracing reveals the dynamic contribution of *Hes1* + cells to the developing and adult pancreas. Development 138, 431–441.

37. Krentz, N.A.J., Lee, M.Y.Y., Xu, E.E., Sproul, S.L.J., Maslova, A., Sasaki, S., and Lynn, F.C. (2018). Single-Cell Transcriptome Profiling of Mouse and hESC-Derived Pancreatic Progenitors. Stem Cell Reports 11, 1551–1564.

38. Langmead, B., Trapnell, C., Pop, M., and Salzberg, S.L. (2009). Ultrafast and memory-efficient alignment of short DNA sequences to the human genome. Genome Biol. 10, R25.

39. Lee, A.-H., Heidtman, K., Hotamisligil, G.S., and Glimcher, L.H. (2011). Dual and opposing roles of the unfolded protein response regulated by IRE1α and XBP1 in proinsulin processing and insulin secretion. PNAS 108, 8885–8890.

40. Lee, C.S., Perreault, N., Brestelli, J.E., and Kaestner, K.H. (2002). Neurogenin 3 is essential for the proper specification of gastric enteroendocrine cells and the maintenance of gastric epithelial cell identity. Genes Dev. 16, 1488–1497.

41. Lee, J., Sugiyama, T., Liu, Y., Wang, J., Gu, X., Lei, J., Markmann, J.F., Miyazaki, S., Miyazaki, J., Szot, G.L., et al. (2013). Expansion and conversion of human pancreatic ductal cells into insulin-secreting endocrine cells. ELife 2.

42. Li, W., Nakanishi, M., Zumsteg, A., Shear, M., Wright, C., Melton, D.A., and Zhou, Q. (2014). In vivo reprogramming of pancreatic acinar cells to three islet endocrine subtypes. Elife 3, e01846.

43. Liu, J., Banerjee, A., Herring, C.A., Attalla, J., Hu, R., Xu, Y., Shao, Q., Simmons, A.J., Dadi, P.K., Wang, S., et al. (2019). Neurog3-Independent Methylation Is the Earliest Detectable Mark Distinguishing Pancreatic Progenitor Identity. Developmental Cell 48, 49–63.e7.

44. Ma, S., Zhang, B., LaFave, L.M., Earl, A.S., Chiang, Z., Hu, Y., Ding, J., Brack, A., Kartha, V.K., Tay, T., et al. (2020). Chromatin Potential Identified by Shared Single-Cell Profiling of RNA and Chromatin. Cell 183, 1103–1116.e20.

45. Masui, T., Swift, G.H., Hale, M.A., Meredith, D.M., Johnson, J.E., and MacDonald, R.J. (2008). Transcriptional Autoregulation Controls Pancreatic Ptf1a Expression during Development and Adulthood. Mol. Cell. Biol. 28, 5458–5468.

46. Mathelier, A., Fornes, O., Arenillas, D.J., Chen, C.-Y., Denay, G., Lee, J., Shi, W., Shyr, C., Tan, G., Worsley-Hunt, R., et al. (2016). JASPAR 2016: a major expansion and update of the open-access database of transcription factor binding profiles. Nucleic Acids Res 44, D110–115.

47. Matys, V., Kel-Margoulis, O.V., Fricke, E., Liebich, I., Land, S., Barre-Dirrie, A., Reuter, I., Chekmenev, D., Krull, M., Hornischer, K., et al. (2006). TRANSFAC and its module TRANSCompel: transcriptional gene regulation in eukaryotes. Nucleic Acids Res 34, D108–110.

48. Maurano, M.T., Humbert, R., Rynes, E., Thurman, R.E., Haugen, E., Wang, H., Reynolds, A.P., Sandstrom, R., Qu, H., Brody, J., et al. (2012). Systematic localization of common disease-associated variation in regulatory DNA. Science 337, 1190–1195.

49. McLean, C.Y., Bristor, D., Hiller, M., Clarke, S.L., Schaar, B.T., Lowe, C.B., Wenger, A.M., and Bejerano, G. (2010). GREAT improves functional interpretation of cis-regulatory regions. Nat Biotech 28, 495–501.

50. Muzumdar, M.D., Tasic, B., Miyamichi, K., Li, L., and Luo, L. (2007). A global double-fluorescent Cre reporter mouse. Genesis 45, 593–605.

51. Newburger, D.E., and Bulyk, M.L. (2009). UniPROBE: an online database of protein binding microarray data on protein-DNA interactions. Nucleic Acids Res 37, D77–82.

52. Olsson, A., Venkatasubramanian, M., Chaudhri, V.K., Aronow, B.J., Salomonis, N., Singh, H., and Grimes, H.L. (2016). Single-cell analysis of mixed-lineage states leading to a binary cell fate choice. Nature 537, 698–702.

53. Petersen, M.B.K., Azad, A., Ingvorsen, C., Hess, K., Hansson, M., Grapin-Botton, A., and Honoré, C. (2017). Single-Cell Gene Expression Analysis of a Human ESC Model of Pancreatic Endocrine Development Reveals Different Paths to β-Cell Differentiation. Stem Cell Reports 9, 1246–1261.

54. Picelli, S., Faridani, O.R., Björklund, Å.K., Winberg, G., Sagasser, S., and Sandberg, R. (2014). Full-length RNA-seq from single cells using Smart-seq2. Nature Protocols 9, 171–181.

55. Qiu, W.-L., Zhang, Y.-W., Feng, Y., Li, L.-C., Yang, L., and Xu, C.-R. (2017a). Deciphering Pancreatic Islet β Cell and α Cell Maturation Pathways and Characteristic Features at the Single-Cell Level. Cell Metabolism 25, 1194–1205.e4.

56. Qiu, X., Mao, Q., Tang, Y., Wang, L., Chawla, R., Pliner, H.A., and Trapnell, C. (2017b). Reversed graph embedding resolves complex single-cell trajectories. Nat Methods 14, 979– 982.

57. Quinlan, A.R., and Hall, I.M. (2010). BEDTools: a flexible suite of utilities for comparing genomic features. Bioinformatics 26, 841–842.

58. Rousseeuw, P.J., Ruts, I., and Tukey, J.W. (1999). The Bagplot: A Bivariate Boxplot. The American Statistician 53, 382–387.

59. Saldanha, A.J. (2004). Java Treeview--extensible visualization of microarray data. Bioinformatics 20, 3246–3248.

60. Scavuzzo, M.A., Hill, M.C., Chmielowiec, J., Yang, D., Teaw, J., Sheng, K., Kong, Y., Bettini, M., Zong, C., Martin, J.F., et al. (2018). Endocrine lineage biases arise in temporally distinct endocrine progenitors during pancreatic morphogenesis. Nature Communications 9, 1–21.

61. Schaffer, A.E., Taylor, B.L., Benthuysen, J.R., Liu, J., Thorel, F., Yuan, W., Jiao, Y., Kaestner, K.H., Herrera, P.L., Magnuson, M.A., et al. (2013). Nkx6.1 Controls a Gene Regulatory Network Required for Establishing and Maintaining Pancreatic Beta Cell Identity. PLoS Genet 9, e1003274.

62. Schwitzgebel, V.M., Scheel, D.W., Conners, J.R., Kalamaras, J., Lee, J.E., Anderson, D.J., Sussel, L., Johnson, J.D., and German, M.S. (2000). Expression of neurogenin3 reveals an islet cell precursor population in the pancreas. Development 127, 3533–3542.

63. Shannon, P., Markiel, A., Ozier, O., Baliga, N.S., Wang, J.T., Ramage, D., Amin, N., Schwikowski, B., and Ideker, T. (2003). Cytoscape: a software environment for integrated models of biomolecular interaction networks. Genome Res 13, 2498–2504.

64. Sharon, N., Chawla, R., Mueller, J., Vanderhooft, J., Whitehorn, L.J., Rosenthal, B., Gürtler, M., Estanboulieh, R.R., Shvartsman, D., Gifford, D.K., et al. (2019). A Peninsular Structure Coordinates Asynchronous Differentiation with Morphogenesis to Generate Pancreatic Islets. Cell 176, 790–804.e13.

65. Siehler, J., Blöchinger, A.K., Meier, M., and Lickert, H. (2021). Engineering islets from stem cells for advanced therapies of diabetes. Nat Rev Drug Discov.

66. Smith, J.P., Corces, M.R., Xu, J., Reuter, V.P., Chang, H.Y., and Sheffield, N.C. (2021). PEPATAC: an optimized pipeline for ATAC-seq data analysis with serial alignments. NAR Genomics and Bioinformatics 3, lqab101.

67. Smith, S.B., Watada, H., and German, M.S. (2004). Neurogenin3 activates the islet differentiation program while repressing its own expression. Mol. Endocrinol. 18, 142–149.

68. Solar, M., Cardalda, C., Houbracken, I., Martín, M., Maestro, M.A., De Medts, N., Xu, X., Grau, V., Heimberg, H., Bouwens, L., et al. (2009). Pancreatic Exocrine Duct Cells Give Rise to Insulin-Producing β Cells during Embryogenesis but Not after Birth. Developmental Cell 17, 849–860.

69. Sugiyama, T., Rodriguez, R.T., McLean, G.W., and Kim, S.K. (2007). Conserved markers of fetal pancreatic epithelium permit prospective isolation of islet progenitor cells by FACS. Proceedings of the National Academy of Sciences 104, 175–180.

70. Tritschler, S., Büttner, M., Fischer, D.S., Lange, M., Bergen, V., Lickert, H., and Theis, F.J. (2019). Concepts and limitations for learning developmental trajectories from single cell genomics. Development 146, dev170506.

71. Veres, A., Faust, A.L., Bushnell, H.L., Engquist, E.N., Kenty, J.H.-R., Harb, G., Poh, Y.-C., Sintov, E., Gürtler, M., Pagliuca, F.W., et al. (2019). Charting cellular identity during human in vitro β-cell differentiation. Nature 569, 368–373.

72. Vierbuchen, T., Ling, E., Cowley, C.J., Couch, C.H., Wang, X., Harmin, D.A., Roberts, C.W.M., and Greenberg, M.E. (2017). AP-1 Transcription Factors and the BAF Complex Mediate Signal-Dependent Enhancer Selection. Molecular Cell 68, 1067–1082.e12.

73. Wang, S., Hecksher-Sorensen, J., Xu, Y., Zhao, A., Dor, Y., Rosenberg, L., Serup, P., and Gu, G.(2008). Myt1 and Ngn3 form a feed-forward expression loop to promote endocrine islet cell differentiation. Dev. Biol. 317, 531–540.

74. Weirauch, M.T., Yang, A., Albu, M., Cote, A.G., Montenegro-Montero, A., Drewe, P., Najafabadi, H.S., Lambert, S.A., Mann, I., Cook, K., et al. (2014). Determination and inference of eukaryotic transcription factor sequence specificity. Cell 158, 1431–1443.

75. White, P., May, C.L., Lamounier, R.N., Brestelli, J.E., and Kaestner, K.H. (2008). Defining Pancreatic Endocrine Precursors and Their Descendants. Diabetes 57, 654–668.

76. Xin, Y., Dominguez Gutierrez, G., Okamoto, H., Kim, J., Lee, A.-H., Adler, C., Ni, M., Yancopoulos, G.D., Murphy, A.J., and Gromada, J. (2018). Pseudotime Ordering of Single Human β-Cells Reveals States of Insulin Production and Unfolded Protein Response. Diabetes 67, 1783–1794.

77. Xu, C.-R., Li, L.-C., Donahue, G., Ying, L., Zhang, Y.-W., Gadue, P., and Zaret, K.S. (2014). Dynamics of genomic H3K27me3 domains and role of EZH2 during pancreatic endocrine specification. The EMBO Journal 33, 2157–2170.

78. Xuan, S., Borok, M.J., Decker, K.J., Battle, M.A., Duncan, S.A., Hale, M.A., Macdonald, R.J., and Sussel, L. (2012). Pancreas-specific deletion of mouse Gata4 and Gata6 causes pancreatic agenesis. J. Clin. Invest. 122, 3516–3528.

79. Yu, X.-X., Qiu, W.-L., Yang, L., Zhang, Y., He, M.-Y., Li, L.-C., and Xu, C.-R. (2019). Defining multistep cell fate decision pathways during pancreatic development at single-cell resolution. The EMBO Journal 38.

80. Zaret, K.S., and Mango, S.E. (2016). Pioneer transcription factors, chromatin dynamics, and cell fate control. Curr. Opin. Genet. Dev. 37, 76–81.

81. Zhang, J., McKenna, L.B., Bogue, C.W., and Kaestner, K.H. (2014). The diabetes gene Hhex maintains -cell differentiation and islet function. Genes & Development 28, 829–834.

